# POGZ suppresses 2C transcriptional program and retrotransposable elements

**DOI:** 10.1101/2022.11.02.514968

**Authors:** XY Sun, TZ Zhang, LX Cheng, W Jiang, YH Sun

## Abstract

The *POGZ* gene has been found frequently mutated in neurodevelopmental disorders (NDDs) such as autism spectrum disorder (ASD) and intellectual disability (ID). We have recently shown that POGZ maintains mouse embryonic stem cells (ESCs) as a chromatin regulator and a transcription factor. However, the exact mechanisms remain unclear. Here we show that POGZ plays important role in the maintenance of ESCs by silencing the *Dux* gene and certain endogenous retroviruses (ERVs). POGZ directly binds to the *Dux* gene and ERVs, and its depletion leads to up-regulation of 2C genes and the repetitive elements such as RLTR9E and IAP (the intracisternal A-type particles), resulting in transition to a 2C-like (2CLC) state and genome instability. POGZ regulates ESC heterochromatin state by association and recruiting TRIM28 and SETDB1, and its loss leads to increased H3K4me3 and H3K27ac, and decreased H3K9me3 at local chromatin. Activation of POGZ-bound ERVs is associated with up-regulation of nearby neural genes. Chimeric transcripts that are initiated within ERVs and spliced to genic exons are highly expressed in *Pogz−/−* ESCs. Our findings establish that POGZ is required for the maintenance of ESCs by repressing *Dux* and silencing ERVs, which may provide important insights into the disease pathology caused by POGZ dysfunction.

**Highlights:**

1. POGZ depletion leads to activation of 2C genes
2. POGZ depletion leads to deregulation of ERVs
3. POGZ directly binds and represses *Dux*
4. POGZ associates with TRIM28/SETDB1 to maintain heterochromatin state to silence ERVs
5. Activation of POGZ-bound ERVs is associated with up-regulation of nearby neural genes

## INTRODUCTION

*POGZ* encodes a transcription factor that contains multiple domains, including a zinc-finger cluster, an HP1-binding motif, a centromere protein B-DB domain, and a transposase-derived DDE domain (Nozawa et al., 2011). POGZ is shown to function as a transcriptional repressor as it is a reader of heterochromatin marks H3K9me2/3 (Vermeulen et al., 2010), is associated with heterochromatin protein 1 (HP1) (Nozawa et al., 2011; Ostapcuk et al., 2018), and inhibits transcription in vitro (Suliman-Lavie et al., 2020). In mice, *Pogz* mRNA is abundantly expressed during early gestation. *Pogz* deletion causes early embryonic lethal with severe phenotypes as early as at E10.5 (Gudmundsdottir et al., 2018; Markenscoff-Papadimitriou et al., 2021), underlying its importance in embryonic neural development. Consistently, conditional deletion of *Pogz* causes dysfunction of mouse brain (Cunniff et al., 2020; Markenscoff-Papadimitriou et al., 2021; Suliman-Lavie et al., 2020). Of note, *POGZ* is one of the top risk genes mutated in neurodevelopmental disorders (NDDs) such as ASD and ID (De Rubeis et al., 2014; Fukai et al., 2015; Matsumura et al., 2020; Stessman et al., 2016; Wang et al., 2016; White et al., 2016; Ye et al., 2015; Zhao et al., 2019). However, the underlying pathology remains obscure.

Under conventional LIF/KSR culture condition, ESCs are heterogeneous and contain a small percentage of cells that not only transiently express ZGA transcripts but also share certain epigenetic feature with the two-cell embryo. These so-called “ 2C-like” ESCs (2CLCs) can be labelled by fluorescent reporters driven by the promoters of the endogenous retrovirus MERVL or *Zscan4* cluster (Zalzman et al. 2010; Macfarlan et al. 2012; Eckersley-Maslin et al. 2016). Compared to ESCs, the totipotent 2CLCs express 2C signature genes such as *MERVL*, *Dux*, *Tcstv3*, and *Zscan4*, exhibit loss of chromocenters, and have a global more open chromatin with increased active histone marks such as H3K27ac and H3K4me3 and decreased repressive histone marks such as H3K9me3 and H4K20me3 (Ishiuchi et al., 2015). An increasing number of 2C/ZGA regulators have been identified, including TRIM28/KAP1 (Rowe et al. 2010; Macfarlan et al. 2011), SETDB1 (Wu et al., 2020), LSD1/*Kdm1a* (Macfarlan et al. 2012), the histone chaperone CAF-1 (Ishiuchi et al. 2015), RIF1 (Li et al., 2017), the LINE1–nucleolin complex (Percharde et al. 2018), LIN28 (Sun et al., 2022), Dapp2/4 (Eckersley-Maslin et al. 2018), and non-coding miRNAs (Yang et al., 2020).

Retrotransposable elements (REs) compose a significant portion of mammalian genomes. During embryogenesis, REs such as endogenous retroviral elements (ERVs) are activated at the 2C stage to promote zygote genome activation (ZGA), and must be effectively silenced thereafter. Previous work has shown that the repression of ERVs is achieved primarily by epigenetic mechanisms involving the formation of heterochromatin and/or DNA methylation downstream of TRIM28/KAP1 and SETDB1 (Karimi et al., 2011; Matusi et al., 2010). However, the upstream regulators and how TRIM28 and SETDB1 are targeted to specific REs remain elusive. Of note, aberrant RE activation in somatic cells is closely linked with various diseases, including neural disorders (Douville et al. 2011; Payer and Burns, 2019; Takahashi et al., 2022).

We have recently shown that POGZ maintains ESCs as a chromatin regulator and a transcription factor (Sun et al., 2022). However, the detailed mechanisms remain unclear. In this work, we found that POGZ is required for repressing 2C genes and ERVs, and its absence induces the emergence of 2CLCs. POGZ associates and recruits TRIM28 and SETDB1 to establish heterochromatin marks on the *Dux* gene and the repetitive elements. Our findings revealed key roles of POGZ in the transcriptional silence of ERVs and *Dux*.

## MATERIALS AND METHODS

### ES Cell Culture

Mouse embryonic stem cells (ESCs) R1 were maintained in Dulbecco’s Modified Eagle Medium (DMEM, BI, 01-052-1ACS) containing 20% knockout serum replacement (KSR, Gibco, 10828028), 1 mM sodium pyruvate (Sigma, S8636), 2 mM L-Glutamine (Sigma, G7513), 1,000 U/mL leukemia inhibitory factor (LIF, Millipore, ESG1107) and penicillin/streptomycin (Gibco, 15140-122) at 37°C with 5% CO2. Cells were routinely propagated by trypsinization and replated every 3-4 days, with a split ratio of 1: 6.

### Generation of *Pogz−/−* ESCs

*Pogz−/−* ESCs were generated by CRISPR/Cas9 technology. Briefly, we designed gRNAs on exon 2 of the *Pogz* gene by using the online website http://crispr.mit.edu/. The gRNA sequence is 5’-CGACCTGTTTATGGAATGTGAGG-3’. The guide sequence oligos were cloned into pSpCas9(BB)-2A-puro containing Cas9 and the sgRNA scaffold (Ran et al., 2013). The plasmid without addition of the gRNA was used as control. The plasmids containing the sgRNA sequence or control plasmids were transfected into ESCs using Neon transfection system according the manufacturer’s instructions. 48 hours later, transfected ESCs were selected with 1 μg/mL puromycin for 4-5 days. Most of cells died within 4 days, and the remaining cells were allowed to proliferate. After 10-15 days selection, the single colony was picked up and passaged into 24 well plates.

DNA from single colonies from the passaged cells was extracted and used for genotyping by PCR. The PCR primer was designed to amplify 200-300 bp on either side of the sgRNA sites. The PCR products were ligated into pGEM-T vector and sequenced to determine the genotype of each single colony. We have successfully generated 3 mutant alleles (1 bp deletion, 7 bp insertion, and 284 bp insertion in exon 2 of the *Pogz* gene). Three homozygous *Pogz*−/− ESC lines have been established: Mutation 1 (Mut1, 1 bp deletion), Mutation 2 (Mut2, 284 bp insertion), and Mutation 3 (Mut3, 7 bp insertion) (Sun et al., 2022).

### Generation of 3×Flag-POGZ restoring *Pogz−/−* mESC Cell Lines

The full-length *Pogz* cDNA (NM_172683.4) was amplified by PCR and then cloned into pCMV-3×Flag vector. The full-length *Pogz* cDNA sequence containing N-terminal 3×Flag sequence was subcloned into pCAG-IRES-Neo vector. To make stable transgenic cells, *Pogz−/−* ESCs were transfected with pCAG-IRES-Neo-3×Flag-*Pogz* vector using Lipofectamine 3000 (Gibco, L3000008). 48 hours later, cells were selected by 500 μg/mL G418 for a week. Cells were expanded and passaged in medium with 200 μg/mL G418. Western Blot assay was performed to confirm the transgenic cell line using Flag antibodies.

### Generation of 2C∷tdTomato reporter mESC Cell Lines

For generation of 2C∷tdTomato reporter (Macfarlan et al., 2012) ESC lines, 1×10^6^ *Pogz+/+* and *Pogz−/−* ESCs were seeded in gelatinized 60 mm dishes, and were transfected with 2C∷tdTomato reporter vector (Miaolingbio, P1701, Wuhan) using Hieff Trans TM Liposomal Transfection reagent (Yeasen, #40802ES03, Shanghai). Cells were selected with 200 μg/mL hygromycin for 7 to 10 days after transfection. The 2C∷tdTomato positive cells were checked by a Leica SP8 laser scanning confocal microscope.

### siRNA knockdown

We used small interfering RNA (siRNA) to knockdown (KD) the *Dux* gene cluster in ESCs. *Dux* siRNAs were perchased from ThermoFisher (#AM16708) and Santa Cruz (sc-144142). siRNAs were transfected into *Pogz−/−* ESCs using Lipofectamine 3000 reagent.After 72h, ESCs were collected for qRT-PCR analysis.

### RNA preparation, RT-qPCR and RNA-seq

Total RNA from mESCs and ESC-derived EBs was extracted with a Total RNA kit (Omega, R6834-01). A total of 1 μg RNA was reverse transcribed into cDNA using the TransScript All-in-One First-Strand cDNA synthesis Supermix (Transgen Biotech, China, AT341). Quantitative real-time PCR (qRT-PCR) was performed using the TransStart® Tip Green qPCR SuperMix (Transgen Biotech, China, AQ-601). The primers used can be found in Supplementary file 1. All qRT-PCR experiments were repeated three times, and the relative gene expression levels were calculated based on the 2^−ΔΔCt^ method. Data were shown as means ± S.D. The Student’s t test was used for the statistical analysis. The significance is indicated as follows: *, *p* < 0.05; **, *p* < 0.01; ***, *p* < 0.001.

For RNA-seq, control and *Pogz* mutant ESCs were collected in tubes pre-loaded with Trizol (Transgen, Beijing). RNAs were quantified by a Nanodrop instrument, and were submitted to BGI Shenzhen (Wuhan, China) for mRNA enrichment, library construction, and sequencing (BGISeq 50SE). RNA-seq was performed in at least two technical repeats and 2 biological replicates. *P*< 0.05 and a Log2 fold change> 1 was deemed to be differentially expressed genes (DEGs).

### Protein extraction and Western blot analysis

For protein extraction, ESCs or HEK293T cells were harvested and lysed in TEN buffer (50 mM Tris-HCl, 150 mM NaCl, 5 mM EDTA, 1% Triton X-100, 0.5% Na-Deoxycholate, supplement with Roche cOmplete Protease Inhibitor). The lysates were quantified by the Bradford method and equal amount of proteins were loaded for Western blot assay. Antibodies used for WB were POGZ (Abcam, ab171934), anti-HP1gamma (Proteintech, 11650-2-AP), Anti-Flag (F1804/F3165, Sigma, 1: 1000), anti-MYC antibody (Transgen Biotech, HT101), anti-H3K9me3 (Abcam, ab176916, 1: 1000), anti-H4K20me3 (Cell Signaling Technology, 5737s, 1: 1000), anti-SETDB1 (Proteintech, 11231-1-AP, 1: 1000) and anti-KAP1 (Proteintech, 15202-1-AP, 1: 1000). Briefly, the proteins were separated by 10% SDS-PAGE and transferred to a PVDF membrane. After blocking with 5% (w/v) non-fat milk for 1 hour at room temperature, the membrane was incubated overnight at 4°C with the primary antibodies. Then the membranes were incubated with a HRP-conjugated goat anti-rabbit IgG (GtxRb-003-DHRPX, ImmunoReagents, 1: 5000), a HRP-linked anti-mouse IgG (7076S, Cell Signaling Technology, 1: 5000) for 1 hour at room temperature. The GE ImageQuant LAS4000 mini luminescent image analyzer was used for photographing. Western blot experiments were repeated at least two times.

### Co-immunoprecipitation assay (co-IP)

Co-IPs were performed with the Dynabeads Protein G (Life Technologies, 10004D) according to the manufacturer’s instructions. Briefly, 1.5 mg Dynabeads was conjugated with antibodies or IgG overnight at 4°C. Antibodies were used are: 8 μg IgG (Proteintech, B900610), or 8 μg anti-POGZ antibody, or 8 μg anti-Flag antibody (Sigma, F3165/F1804), or 8 μg anti-Myc antibody (Abbkine, A02040) or 8 μg anti-Myc antibody (Transgen Biotech, HT101). The next day, total cell lysates and the antibody-conjugated Dynabeads were incubated overnight at 4°C with shaking. After three-times washing with PBS containing 0.1% Tween, the beads were boiled at 95°C for 5 minutes with the 6×Protein loading buffer and the supernatant was collected for future WB analysis.

The full-length Setdb1 cDNA (NM_001163641.1) was amplified by PCR and then cloned into pCS2-Myc vector. For transfection, HEK293T cells were seeded in 10 cm dishes and contransfected with pCMV-3×Flag-*Pogz* and pCS2-Myc-*Setdb1* vector using Liposomal Transfection reagent (Yeasen, 40802ES03) for 24 h. Cells were harvested for Co-immunoprecipitation assay (co-IP) and Western blot analysis.

### Flow cytometry analysis and cell sorting

The 2C∷tdTomato reporter mESCs were dissociated into single cells with 0.25% trypsin. Then 2C∷tdTomato positive cells and negative cells were sorted by the BD-AriaIII flow cytometry system in the R&D Division of Public Technology and Service of IHB (preformed by Dr. Wang Yan). Data were analyzed by FlowJo 10.5.0 sofeware (TreeStar, OR, USA).

### Luciferase reported assays

The *Dux* promoter was amplified by PCR with PCR mix (Yeasen, 10149ES03) and then cloned into pGL4.10[luc2] (Addgene, E6651) vector. For luciferase reporter assay in HEK293T, 1×10^5^ cells were seeded in 24-well plates in DMEM medium containing 10% FBS (Transgen Biotech, FS101-02). The cells were transfected with the indicated vectors using Hieff Trans TM Liposomal Transfection reagent (Yeasen, 40802ES03).

For luciferase reporter assay in ESCs, the vectors were transfected using the Neon Transfection System as described previously (Sun et al., 2020). Rellina was used as an internal control. The luciferase activity was measured with Dual-luciferase Reporter Assay System (Promega, Madison, E1910). Experiments were repeated three times. Data were represented as mean± SEM based on three independent measurements. Statistical significance was calculated by two-tailed unpaired or paired Student’s t-test.

### Chromatin Immunoprecipitation (ChIP) and ChIP-seq

ChIP experiments were performed according to the Agilent Mammalian ChIP-on-chip manual as described (Sun et al., 2022). Briefly, 1× 10^8^ ES cells were fixed with 1% formaldehyde for 10 minutes at room temperature. Then the reactions were stopped by 0.125 M Glycine for 5 min with rotating. The fixed chromatin were sonicated to an average of 200-500bp (for ChIP-Seq) or 500-1,000bp (for ChIP-qPCR) using the S2 Covaris Sonication System (USA). For ChIP-seq, chromatin was sheared for 15 min with Peak power:135; Duty factor: 5.0; Cycles/Burst 200. Then Triton X-100 was added to the sheared chromatin solutions to a final concentration of 0.1%. After centrifugation, 50 μl of supernatants were saved as input. The remainder of the chromatin solution was incubated with Dynabeads previously coupled with 10 μg ChIP grade antibodies (POGZ, Abcam, ab167408;Flag, Sigma, F1804/F3165; H3K4me3, Abcam, ab1012; H3K27ac, Millipore, MABE647; H3K9me3, Abcam, ab176916; H4K20me3, Cell Signaling Technology, 5737s; TRIM28, Proteintech, 15202-1-AP; SETDB1, Proteintech, 11231-1-AP) overnight at 4°C with rotation. Next day, after 7 times washing with the wash buffer, the complexes were reverse cross-linked overnight at 65°C. DNAs were extracted by hydroxybenzene-chloroform-isoamyl alcohol and purified by a Phase Lock Gel (Tiangen, China). For ChIP-seq, the ChIPed DNA were dissolved in 15 μl distilled water. Library constructions and deep sequencing were done by the BGI Shenzhen (Wuhan, China). The ChIP-seq experiments were repeated three times for each antibody (using POGZ antibodies in control ESCs, and using FLAG antibodies in FLAG-tagged POGZ restoring *Pogz−/−* ESCs).

For ChIP-PCR, the ChIPed DNA were dissolved in 100 μl distilled water. Quantitative real-time PCR (qRT-PCR) was performed using a Bio-Rad qPCR instrument. The enrichment was calculated relative to the amount of input. All experiments were repeated at least three times. The relative gene expression levels were calculated based on the 2^− Δ Δ Ct^ method. Data were shown as means ± S.D. The Student’s t test was used for the statistical analysis. The significance is indicated as follows: *, *P*< 0.05; **, *P*< 0.01; ***, *P*< 0.001.

### CUT&Tag

The Cleavage Under Targets and Tagmentation (CUT&Tag) experiments were performed using FLAG antibodies (Flag, Sigma, F1804/F3165) in 3× Flag-POGZ restoring *Pogz−/−* mESCs by the Yingzi and Frasergen Companies (Wuhan, China) (Kaya-Okur et al., 2019). The detailed methods have been recently described (Sun et al., 2022).

### Quantification and statistical analysis

Data are presented as mean values ±SD unless otherwise stated. Data were analyzed using Student’s t-test analysis. Error bars represent s.e.m. Differences in means were statistically significant when *P*< 0.05. Significant levels are: \**P*< 0.05; \*\**P*< 0.01, and \*\*\**P*< 0.001.

### Bioinformatics

#### RNA-seq analysis

The RNA-seq data were aligned to the GRCm39 reference genome using HISTA2. Then, raw counts of all protein coding genes was generate by FeatureCounts. Raw counts was normalized to TPM (transcript per million). DESeq2 was performed to calculate differentially expressed genes with abs|log2(fold change)|> 1 and *P*-adj < 0.05.

For retrotransposon elements analysis, only the best alignments were kept while multi-mapped reads were randomly retained once. CPM (counts per million) of REs were calculated. Bedtools was used for the counting of individual sites and only uniquely mapped reads were kept for individual LTR.

#### ChIP-seq and CUT&Tag analyses

For ChIP-seq analysis, the reads were aligned to the GRCm39 reference genome using bowtie2 with the parameters: −p 64 – very-sensitive – end-to-end – no-unal. PCR duplicates were removed using Picard MarkDuplicates. Peaks were called using MACS2. BigWig files were generated using deeptools with the RPKM normalization method.

For CUT&Tag analysis, technical sequences and sequencing adapters were trimmed using the Trimmomatic tool. Raw data were evaluated using FastQC. Each sample was aligned in Bowtie2 with the options: -I 10 -X 1000 --dovetail --no-unal --very-sensitive-local --no-mixed --no-discordant. Non-aligning and multiple mappers were filtered out by Picard tools. Peaks were called using MACS2 (version 2.1.1) with the parameters: -f BAMPE -B --call-summits --keep-dup all. BigWig files were generated using deeptools with the RPKM normalization method.

For retrotransposon elements analysis, the annotations of transposon elements were obtained from UCSC Genome Browser RepeatMasker. We filtered out annotations < 300 bp, only the best alignments were kept while multi-mapped reads were randomly retained once, all REs were treated as meta-gene. For RE enrichment analysis, we calculated the total counts of each family element in the annotations of REs, then calculated the IP/input log2(fold change) ratio.

#### ATAC-seq Analysis

Paired-end reads were aligned using Bowtie2 using default parameters. Only uniquely mapping reads were kept for further analysis. These uniquely mapping reads were used to generate bigwig genome coverage files similar to ChIP–seq. Heat maps were generated using deeptools2. For the meta-profiles, the average fragment count per 10-bp bin was normalized to the mean fragment count in the first and last five bins, which ensures that the background signal is set to one for all experiments. Merged ATAC-seq datasets were used to extract signal corresponding to nucleosome occupancy information with NucleoATAC.

## RESULTS

### POGZ depletion induces 2CLCs

We have recently generated three *Pogz−/−* ESC lines (Mut1-3), by CRISPR/Cas9 technology (Sun et al., 2022). Similar phenotypes, including genome instability, growth and self-renewal/proliferation defects, were observed in all three mutant ESCs (Nozawa et al., 2010). To investigate the mechanism by which POGZ maintains ESCs, RNA sequencing (RNA-seq) was performed for control and early passage *Pogz−/−* ESCs. There was little changes of pluripotency and three germ layer marker genes, suggesting that acute POGZ depletion does not induce precocious differentiation. Interestingly, we found that 2C-specific genes such as the *Zscan4* cluster (such as *Zscan4b/c/d/f*), *Zfp352*, MERVL, *Fth17ls* and *Tcstv3*, were markedly activated in *Pogz−/−* ESCs (Figure 1A; Supplementary Figure 1A).

**Figure 1.**
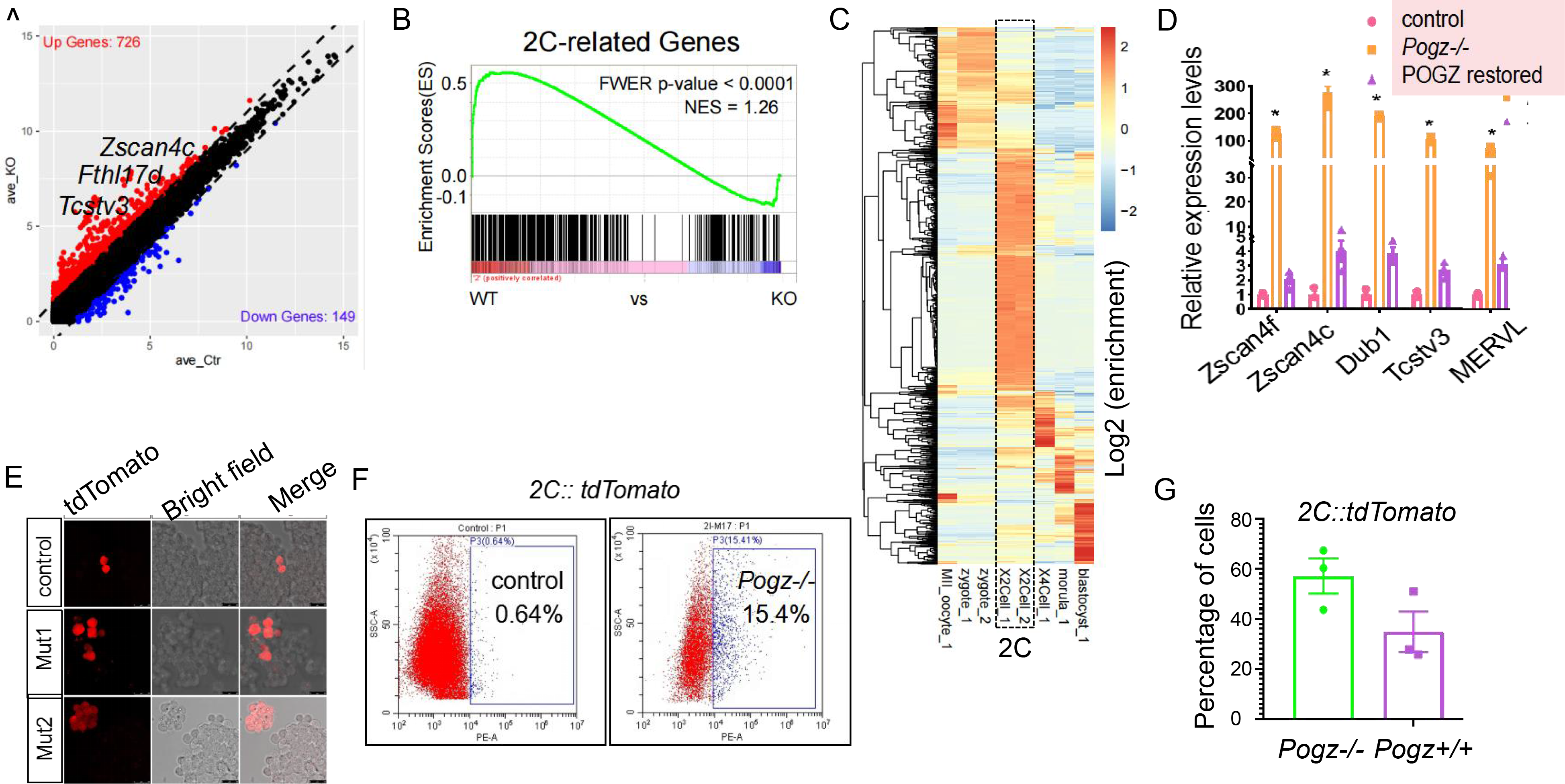
Loss of POGZ induces 2C-like cells. (A) Vacano plot showing the DEGs between control and *Pogz−/−* ESCs. (B) Gene set enrichment analysis showing the up-regulation of 2C genes upon loss of POGZ. (C) Heat map showing the dynamic expression of up-regulated DEGs during mouse early embryogenesis, based the published RNA-seq data (Wang, et. al., 2018). Note: roughly half of the DEGs are 2C-stage specific (black dashed box). (D) qRT-PCR analysis of selected 2C genes in control, *Pogz*−/−, and Flag-POGZ restored *Pogz−/−* ESCs. (E) Representative images showing the 2C∷tdTomato positive cells in control, Mut1 and Mut2 ESCs. (F) Flow results showing the percentage of 2C∷tdTomato positive cells in control and Mut1 ESCs. (G) Percentage of 2C-tdTomato positive cells from *Pogz+/+* and *Pogz−/−* transgenic ESCs after sorting and replating after 3 days. The qRT-PCR were repeated three times.

The activation of 2C genes prompted us to perform gene set enrichment analysis (GSEA) of the up-regulated differentially expressed genes (DEGs), and the results showed that they were significantly enriched in the 2C gene signature (Figure 1B). Consistently, the heat map analysis showed that the up-regulated DEGs were highly enriched for 2C stage and zygotic genome activation (ZGA) genes (Figure 1C; Supplementary Figure 1B) (Wang et al., 2018). By qRT-PCR, we confirmed that the selected 2C genes, including *MERVL*, *Zscan4*, *Tcstv3*, *Dub1*, *Usp17ls, Todpoz3*, *Sp110*, *Gm4340*, and *Zfp352*, were up-regulated in *Pogz−/−* ESCs, which can be rescued by reintroducing the Flag-tagged POGZ (Figure 1D; Supplementary Figure 1C). These observations suggested that loss of POGZ induces the emergence of the 2CLCs.

The 2CLCs can be labelled using fluorescent reporters driven by the promoters of the endogenous retrovirus MERVL or *Zscan4* (Zalzman et al. 2010; Macfarlan et al. 2012). We established 2C-tdTomato transgenic ESCs, where the red fluorescenct protein tandem dimeric Tomato (tdTomato) is driven by a MERVL-LTR, a member of the MERVL family of retroviral elements. Under the fluorescence microscope, a marked increase of 2C-tdTomato positive cells were observed in *Pogz−/−* transgenic ESCs, compared to *Pogz+/+* transgenic ESCs (Figure 1E). Flow cytometry assay showed that a small percentage (0.64%) of *Pogz+/+* transgenic ESCs were tdTomato positive, whereas in *Pogz−/−* transgenic ESCs, the percentage of tdTomato positive cells was increased to 15.4% (Figure 1F).

Next, we sorted 2C-tdTomato positive cells from *Pogz+/+* and *Pogz−/−* transgenic ESCs, and replated them into 24-well plates. FACS-sorted cells fluctuate, which is consistent with the fluctuating nature of 2CLC population in ESCs. Remarkably, 3 days after FACS sorting, 58% of the sorted *Pogz−/−* cells still remained tdTomato positive, whereas only 34% of the sorted *Pogz+/+* transgenic cells were tdTomato positive (Figure 1G). This observation suggested that POGZ restrains 2CLC state in ESCs.

### POGZ transcriptionally represses ERVs

The 2C-like state in ESCs is featured by activation of specific REs such as MERVL and its LTR MT2-Mm (Macfarlan et al., 2012). POGZ contains a transposase-derived DDE domain and a HP1-binding motif (Yuan and Wessler, 2011; Nozawa et al., 2011). In physiological conditions, POGZ associates with heterochromatin proteins 1 (HP1) which is known to be involved in RE regulation (Rowe et al., 2010). We therefore asked whether loss of POGZ leads to deregulation of REs. Analysis of the RNA-seq data showed that this was indeed the case. Approximately 50 endogenous retrovirus (ERVs), belonging to class I (such as MuLV-int/RLTR4_Mm, MMERGLN-int, LTRIS2) and class II (such as IAP elements, RLTR6/9E) types, were significantly deregulated (fold change> 2; *P*< 0.05) (Figure 2A; Supplementary Figure 2A). POGZ depletion primarily resulted in activation of ERVs that are normally expressed in 2C-and later developmental stages of early mouse embryos (Figure 2B), which suggested that POGZ is persistently required to silence REs throughout embryogenesis. The qRT-PCR analysis of the selected REs confirmed the RNA-seq results (Figure 2C; Supplementary Figure 2B). Importantly, reintroducing the Flag-tagged POGZ rescued the expression of the selected repetitive elements.

**Figure 2.**
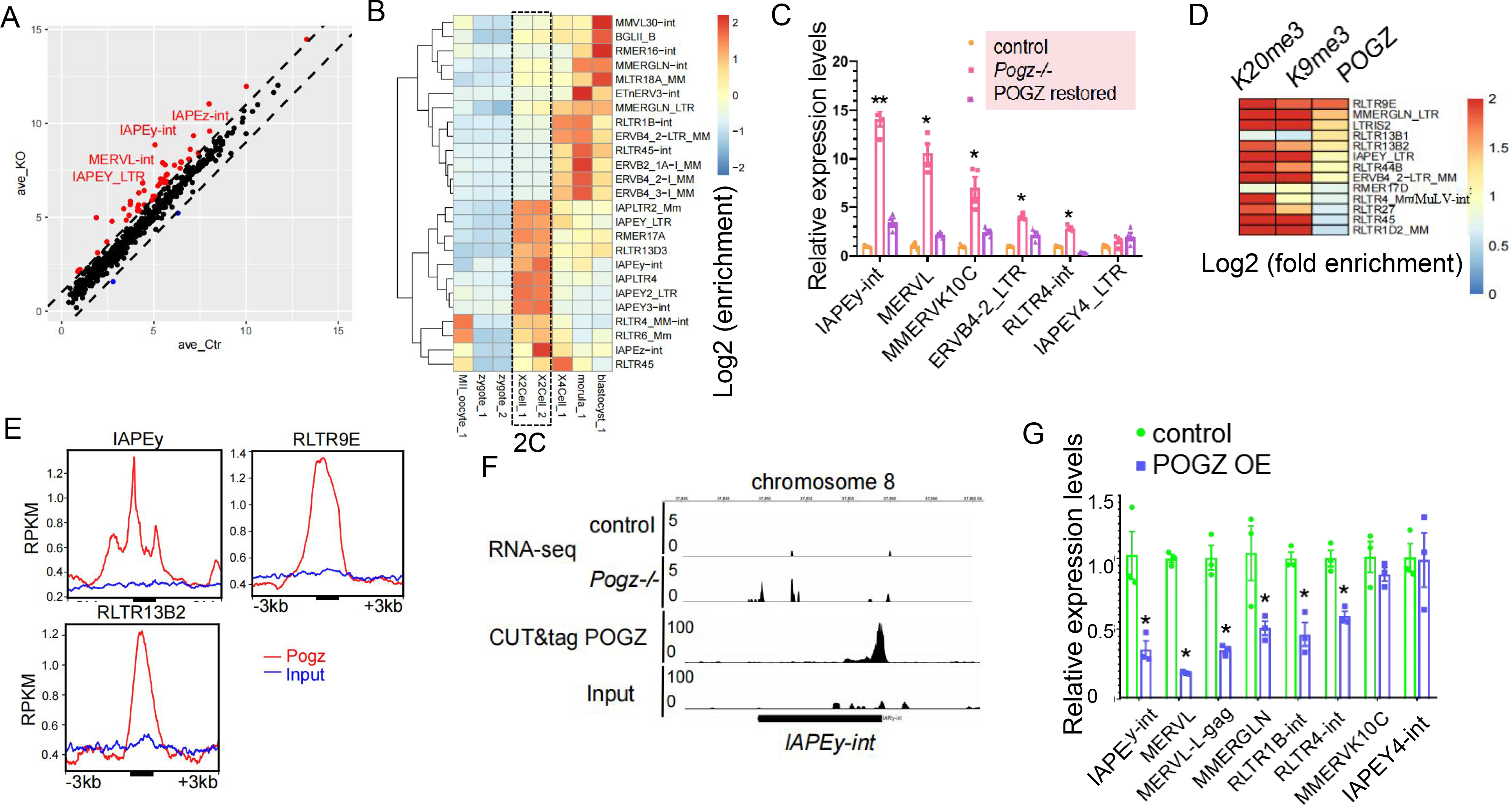
POGZ transcriptionally represses ERVs. (A) Volcano plot showing the differentially expressed REs between control and *Pogz−/−* ESCs. (B) Heat map showing the dynamic expression of top 25 up-regulated REs during early mouse embryogenesis. REs that are expressed in 2C stage of mouse embryos were highlighted (black dashed box). (C) qRT-PCR analysis of selected ERVs in control, *Pogz*−/−, and Flag-POGZ restored *Pogz−/−* ESCs. IAPEY4_LTR, which was not bound by POGZ, was served as a negative control. (D) Heat map showing the enrichment of H3K9me3 and H4K20me3 on the POGZ-bound ERVs. (E) RPKM showing the enrichment of POGZ on the indicated ERVs. RPKM: reads per kilobase of transcripts, per million mapped reads. (F) Genomic views of RNA-seq and CUT&Tag data for IAPEy-int. (G) qRT-PCR analysis of selected ERVs in control and *Pogz* overexpressing ESCs. MMERVK10c and IAPEY4-int (free of POGZ enrichment) were served as negative controls. The qRT-PCR experiments were repeated three times.

To investigate how POGZ is involved in the regulation of ERVs, we examined the genome-wide binding profile of POGZ by performing ChIP-seq (chromatin immunoprecipitation followed sequencing) and CUT&Tag experiments (Kaya-Okur, et al. 2019). Of the 50 deregulated ERV families, about 10 showed significant POGZ enrichment (Figure 2D-F), with RLTR9E being the top one, which suggested that POGZ controls ERVs directly and indirectly. Interestingly, POGZ-bound ERVs are among the youngest murine TE elements and have retained the ability to transpose (Stocking and Kozak, 2008), implying the functional specificity of POGZ.

On the other hand, we overexpressed POGZ in ESCs. Compared to the control ESCs, POGZ overexpression decreased the expression of POGZ-bound ERVs, but had little effects on ERVs free of POGZ binding (Figure 2G). Based on the loss- and gain-of-function data, we concluded that POGZ plays key role in transcriptional silence of ERVs, in particular the recently evolved ERVs.

### POGZ negatively regulates *Dux*

Next, we investigated the mechanisms by which POGZ depletion induces 2CLCs. Induction of 2C-like state can be due to up-regulation of 2CLC positive regulators or down-regulation of its repressors. We found that genes encoding known repressors of the 2C-like state, including *Lin28a*, *Trim28, Kdm1a, Ncl*, and *Chaf1* were normally expressed (Eckersley-Maslin et al., 2018; Percharde et al., 2018; Sun et al., 2022). We thus turned to the positive regulators of the 2CLCs or ZGA, including MERVL, *Dux*, *Zscan4* and *Dppa2/4*. *Dppa2/4* were initially excluded as they were normally expressed in *Pogz−/−* ESCs (Supplementary Figure 3A). Although the *Zscan4* cluster and MERVL were markedly up-regulated in *Pogz−/−* ESCs, the ChIP-seq/CUT&Tag data did not show the direct binding of POGZ (Supplementary Figure 3B).

**Figure 3.**
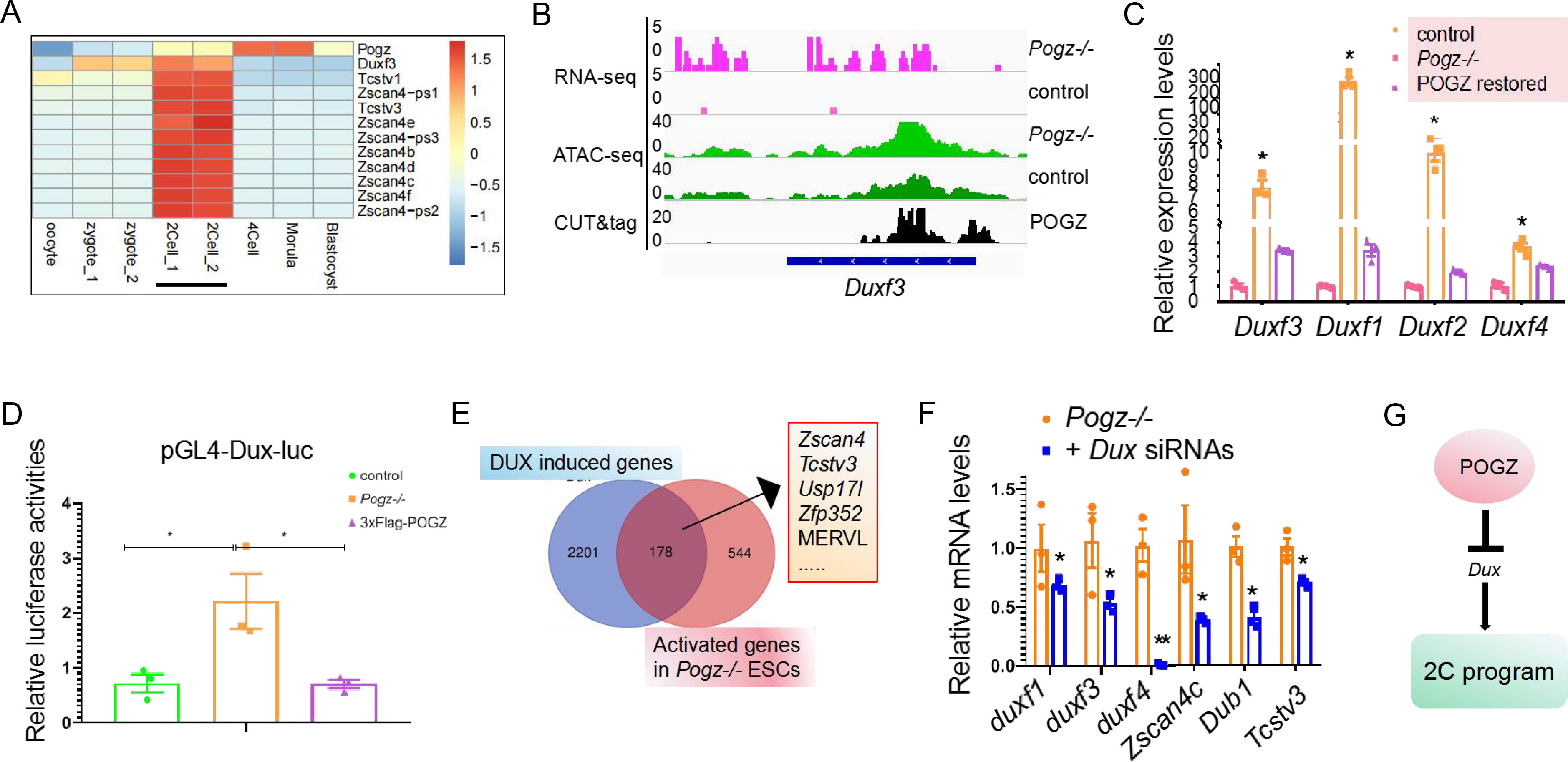
POGZ controls ESCs by repressing *Dux*. (A) Heat map showing the expression of *Pogz*, *Duxf3* and the indicated 2C genes during mouse embryogenesis from published RNA-seq data (Wang, et. al., 2018). (B) Genomic views of RNA-seq, ATAC-seq and CUT&Tag data for the *Duxf3* gene. (C) qRT-PCR analysis of the *Dux* cluster genes in control, *Pogz*−/−, and Flag-POGZ restored *Pogz−/−* ESCs. *Duxf3*: *Dux*. (D) Relative luciferase activities of pGL4-Dux-luc in control, *Pogz*−/−, and Flag-POGZ restored *Pogz−/−* ESCs. (E) Pie chart showing the overlap of DUX overexpression and POGZ depletion induced genes. (F) qRT-PCR analysis of the selected 2C genes in *Pogz−/−* ESCs and *Dux* knockdown *Pogz−/−* ESCs. (G) Cartoon showing that POGZ represses 2C program via transcriptionally silencing *Dux*. The qRT-PCR and luciferase experiments were repeated three times.

*Dux*, which encodes a double homeobox transcription factor, has been identified and considered as a master inducer of 2CLCs (Guo et al., 2019). DUX can bind and robustly activate 2C stage-specific ZGA transcripts, and convert ESCs to a 2C-like state with unique features that resembles the 2C embryos (Hendrickson et al., 2017; Whiddon et al., 2017). Analysis of expression profile of early mouse embryos showed that *Pogz* expression was negatively correlated with *Dux* expression, exhibiting its peak immediately after 2C embryo stage (Figure 3A). This expression pattern suggested that endogenous POGZ is required to suppress the 2C/ZGA genes, which is important for transition from 2C to the next developmental stage. Analysis of Flag-POGZ CUT&Tag data showed that POGZ was highly enriched at the proximal promoter and gene body of the *Dux* gene (Figure 3B). In the absence of POGZ, *Dux* was derepressed, accompanied by an increase in chromatin accessibility as revealed by ATAC-seq (Assay for Transposase Accessible Chromatin with high-throughput sequencing) (Figure 3B). The qRT-PCR analysis confirmed the up-regulation of the *Dux* cluster, which can be largely rescued by reintroducing the Flag-tagged POGZ into *Pogz−/−* ESCs (Figure 3C).

Next, putative DNA binding sites for POGZ were identified by HOMER analysis, with the most highly enriched motif being CCAGGCT (Supplementary Figure 3C). Examination of promoter proximal of the *Dux* gene revealed the existence of such DNA sequences (Supplementary Figure 3D), which prompted us to ask whether POGZ directly represses *Dux* by binding to it. A pGL4-Dux luciferase reporter construct under control by a 2 kb *Dux* promoter was made, and transfected into control and *Pogz−/−* ESCs. The luciferase assay results showed that *Dux* promoter activities were higher in *Pogz-*/-ESCs than control ESCs. Importantly, reintroducing the Flag-tagged POGZ in *Pogz-*/-ESCs restored the luciferase activities (Figure 3D). Moreover, addition of POGZ decreased the pGL4-Dux luciferase activities in HEK293T cells (Supplementary Figure 3E). When CCAGGCT was mutated to CCAaaCT, addition of POGZ had no effects (Supplementary Figure 3F). These results showed that POGZ represses the *Dux* gene by directly binding to it.

DUX overexpression in ESCs was shown to be sufficient to induce 2C or cleavage-stage genes (Hendrickson et al., 2017). We speculated that *Dux* is downstream and mediates POGZ in the induction of 2CLCs in *Pogz−/−* ESCs. To this end, we compared our RNA-seq data with the previous one after *Dux* overexpression, and found that about 180 POGZ depletion induced genes (fold change >2) overlapped with DUX induced genes, which were enriched 2C signature genes such as *Zscan4* and MERVL (Figure 3E). Next, the *Dux* cluster was knocked down in *Pogz−/−* ESCs with siRNAs. *Dux* knockdown rescued the expression of the selected 2C genes, as well as ESC proliferation defects (Figure 3F; Supplementary Figure 3G). Based on these data, we concluded that POGZ maintains ESCs at least partially through repressing *Dux*, the key driver that initiates 2C-like transcription (Figure 3G).

### POGZ controls a subset of ERVs independent of Dux

POGZ target gene *Dux* is known to directly activate 2C-specific repetitive elements such as MERVL (Supplementary Figure 4A) (Hendrickson et al., 2017). It was possible that up-regulation of ERVs in *Pogz−/−* ESCs is a part of 2CLC induction due to increased expression of *Dux*. Alternatively, POGZ may regulate a subset of ERVs independent of *Dux*. To investigate this, we sorted tdTomato positive and negative cells from *Pogz−/−* transgenic ESCs, and examined the expression of the 2C genes and the selected REs (Figure 4A). As expected, the expression of 2C specific genes such as MERVL, *Zscan4* and *Dux* were markedly higher in tdTomato positive cells than in tdTomato negative cells. POGZ-bound ERVs such as IAPEy and MMERGLN were expressed at higher levels in both tdTomato^+^ and negative cells in *Pogz−/−* transgenic ESCs, compared to tdTomato^−^ *Pogz+/+* ESCs (Figure 4A). These observations suggested that POGZ-bound ERVs were up-regulated in all *Pogz−/−* ESCs, which was independent of 2C program. Consistently, when *Dux* was knockdown in *Pogz−/−* ESCs, *Dux* target MERVL was down-regulated as expected, POGZ-bound ERVs, however, remained unchanged (Figure 4B).

**Figure 4.**
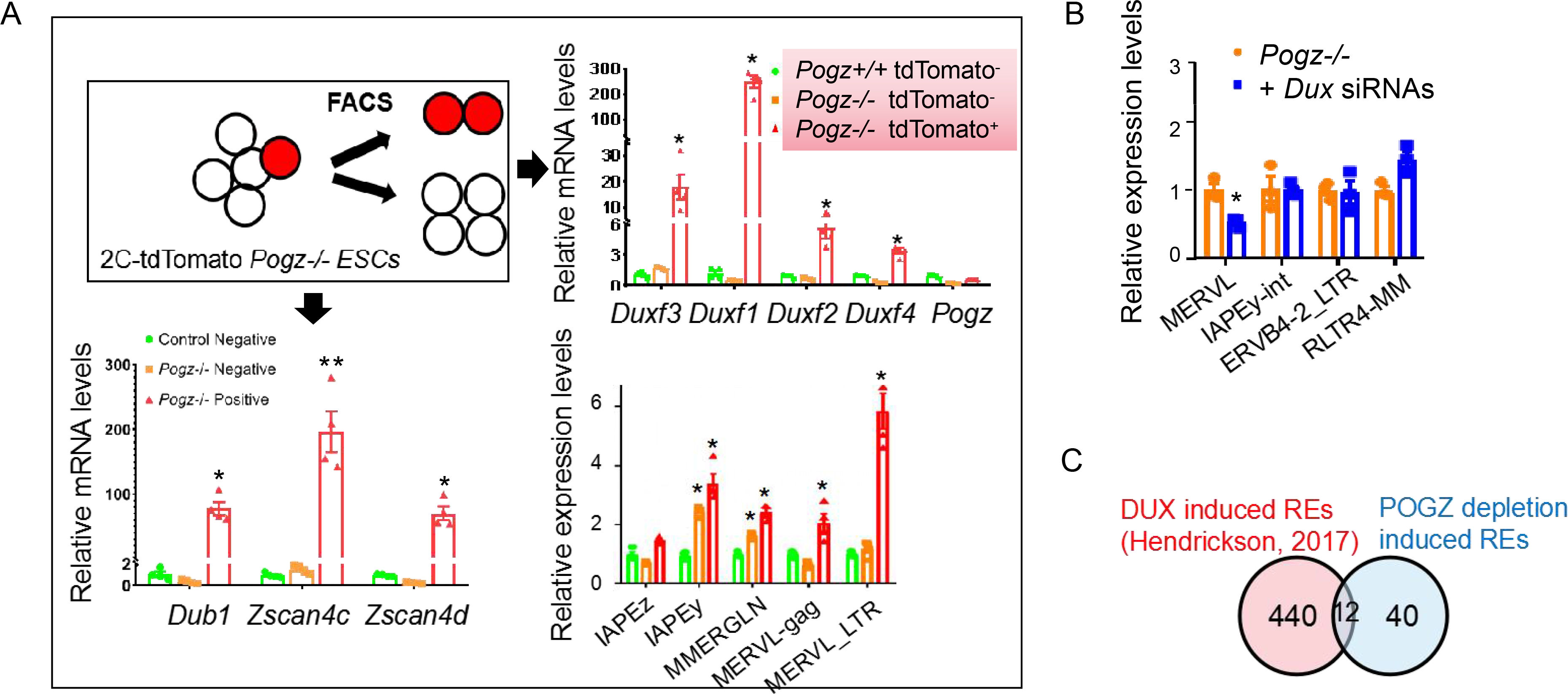
POGZ controls ERVs independent of Dux. (A) qRT-PCR analysis of the indicated ERVs and 2C genes in sorted tdTomato positive and negative cells from *Pogz−/−* transgenic ESCs. Black box showing the experimental design. (B) siRNA KD of *Dux* cluster in *Pogz−/−* ESCs failed to rescue the expression of the indicated POGZ-bound ERVs, except MERVL which is the known target of DUX but not POGZ. (C) Pie chart showing the overlapping REs induced by DUX overexpression and POGZ depletion.

In fact, other observations also supported the above notion. First, many of the POGZ-depletion induced ERVs are not confined to 2C/ZGA stage (Supplementary Figure 2B). Second, there was only a small number of overlapping REs that are induced by POGZ depletion and DUX overexpression (Figure 4C). Third, there was only a few shared ERV families that were activated in the absence of Dux and POGZ (Supplementary Figure 4B-C).

### POGZ maintains local heterochromatin state

Although the majority of POGZ ChIP-seq peaks are localized to gene promoter and enhancer regions decorated with H3K27ac, approximately 8% are localized to heterochromatin regions decorated with H3K9me3/H4K20me3 (Markenscoff-Papadimitriou et al., 2021; Sun et al., 2022). We reasoned that POGZ may regulate ERVs and *Dux* by modulating heterochromatin histone marks, especially considering that the genomic locations of REs and *Dux* are within heterochromatin regions (Liu et al., 2021; Xu et al., 2021). We therefore performed H3K9me3 and H4K20me3 ChIP-seq experiments in control and *Pogz−/−* ESCs.

At the global level, only a small portion of POGZ peaks were overlapped with those of H3K9me3 and H4K20me3 (Supplementary Figure 5A), which was in line with the previous work (Markenscoff-Papadimitriou et al., 2021). Nevertheless, H3K9me3 and H4K20me3 ChIP-seq signals were sightly reduced in the absence of POGZ, which was confirmed by Western blot analysis (Figure 5A). Specifically, POGZ and H3K9me3/H4K20me3 co-occupied the *Dux* gene and ERVs such as IAPEy and RLTR9E (Figure 2D; Figure 5B-C). Importantly, H4K20me3 and H3K9me3 ChIP-seq signals at POGZ-bound IAPEy and *Dux* were markedly reduced in the absence of POGZ (Figure 5C-E; Supplementary Figure 5B).

**Figure 5.**
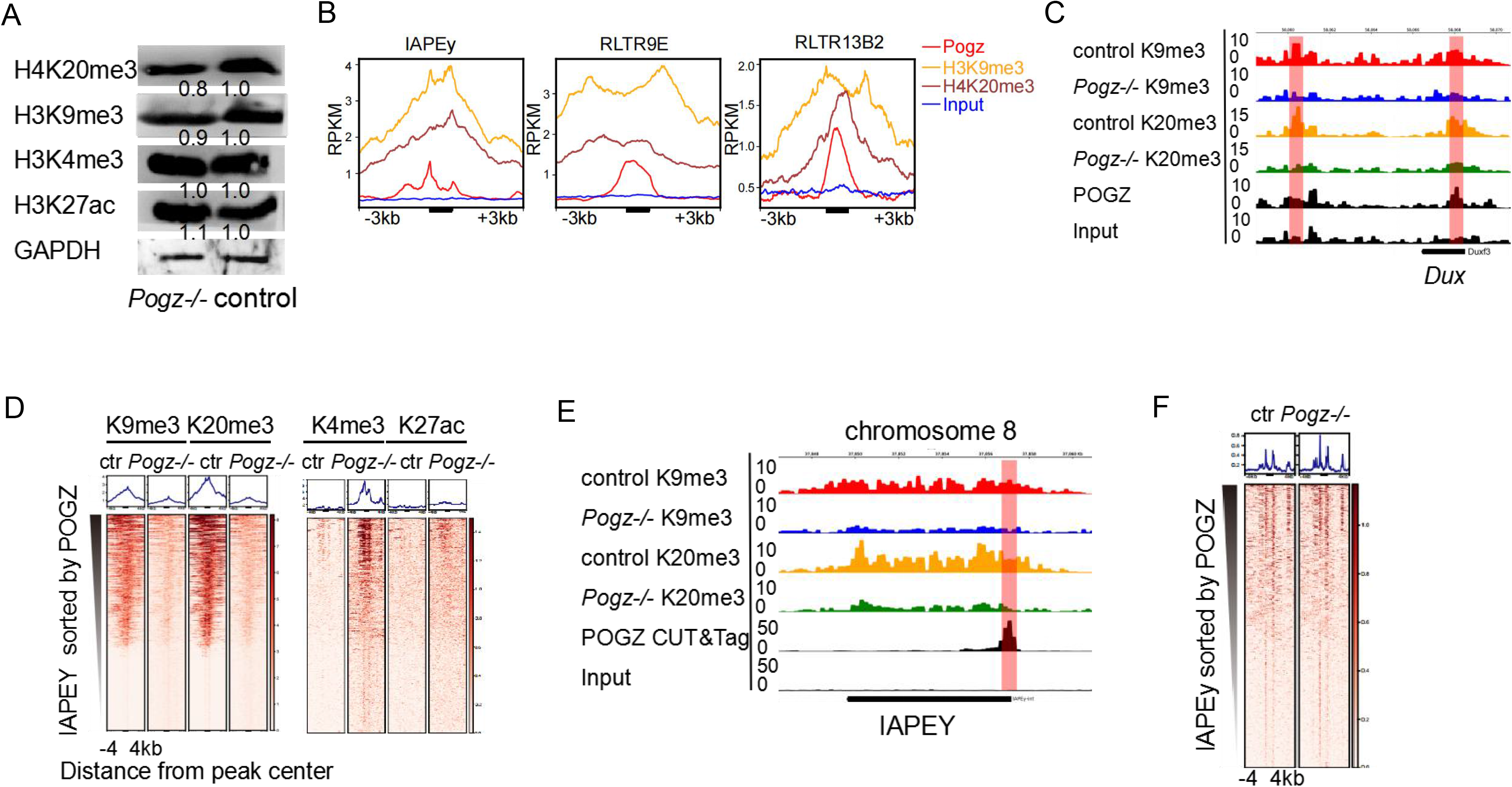
POGZ modulates heterochromatin formation. (A) Western blot image showing the expression of the indicated histone marks in control and *Pogz−/−* ESCs. (B) Read count tag density pileups of the indicated ChIP profiles for the indicated ERVs. RPKM: reads per kilobase of transcript, per million mapped reads. (C) Genomic views of ChIP-seq and CUT&Tag data of POGZ, H3K9me3 and H4K20me3 on *Dux* in control and *Pogz−/−* ESCs. (D) Heat map showing the enrichment of H3K9me3, H4k20me3, H3k4me3 and H3K27ac on POGZ-bound IAPEy in control and *Pogz−/−* ESCs, sorted by POGZ CUT&Tag signals. (E) Genomic views of ChIP-seq and CUT&Tag data of POGZ, H3K9me3 and H4K20me3 on IAPEy in control and *Pogz−/−* ESCs. (F) Heat map showing the ATAC-seq signals on IAPEy in control and *Pogz−/−* ESCs. Western blot experiments were repeated three times.

We also analyzed our previously generated ChIP-seq data of active histone marks such as H3K4me3 and H3K27ac (Sun et al., 2022). When examining the ChIP-seq signals on POGZ-bound ERVs such as IAPEy, a marked increase of H3K4me3 and a moderate increase of H3K27ac were observed in *Pogz−/−* ESCs (Figure 5D; Supplementary Figure 5C). Consistently, chromatin became more open on POGZ-bound ERVs such as IAPEy (Figure 5F). Taken together, we concluded that POGZ is required to establish epigenetic silencing of *Dux* and ERVs, and its loss leads to increased H3K4 trimethylation and H3K27 acetylation, and decreased H3K9 trimethylation, resulting in activation of *Dux* and the repetitive elements.

### POGZ associates with and recruits TRIM28/SETDB1

Previous work have shown that TRIM28 acts upstream of SETDB1 and HP1 to silence ERVs (Matsui et al., 2010), and that the majority of SETDB1-dependent H3K9me3 peaks reside on REs (Wu et al., 2020). As our IP-Mass Spec identified TRIM28 as a potential POGZ interactor (Sun et al., 2022), we reasoned that POGZ may maintain epigenetic silencing of *Dux* and ERVs by association with TRIM28 and/or SETDB1. To investigate this, we first compared POGZ CUT&Tag data with the previously published SETDB1 and TRIM28 ChIP-seq data, and found that POGZ, TRIM28 and SETDB1 shared approximately 8-10% of its binding sites (Supplementary Figure 6A-B). Importantly, the three factors co-occupied POGZ-bound ERVs and *Dux* (Figure 6A-B).

**Figure 6.**
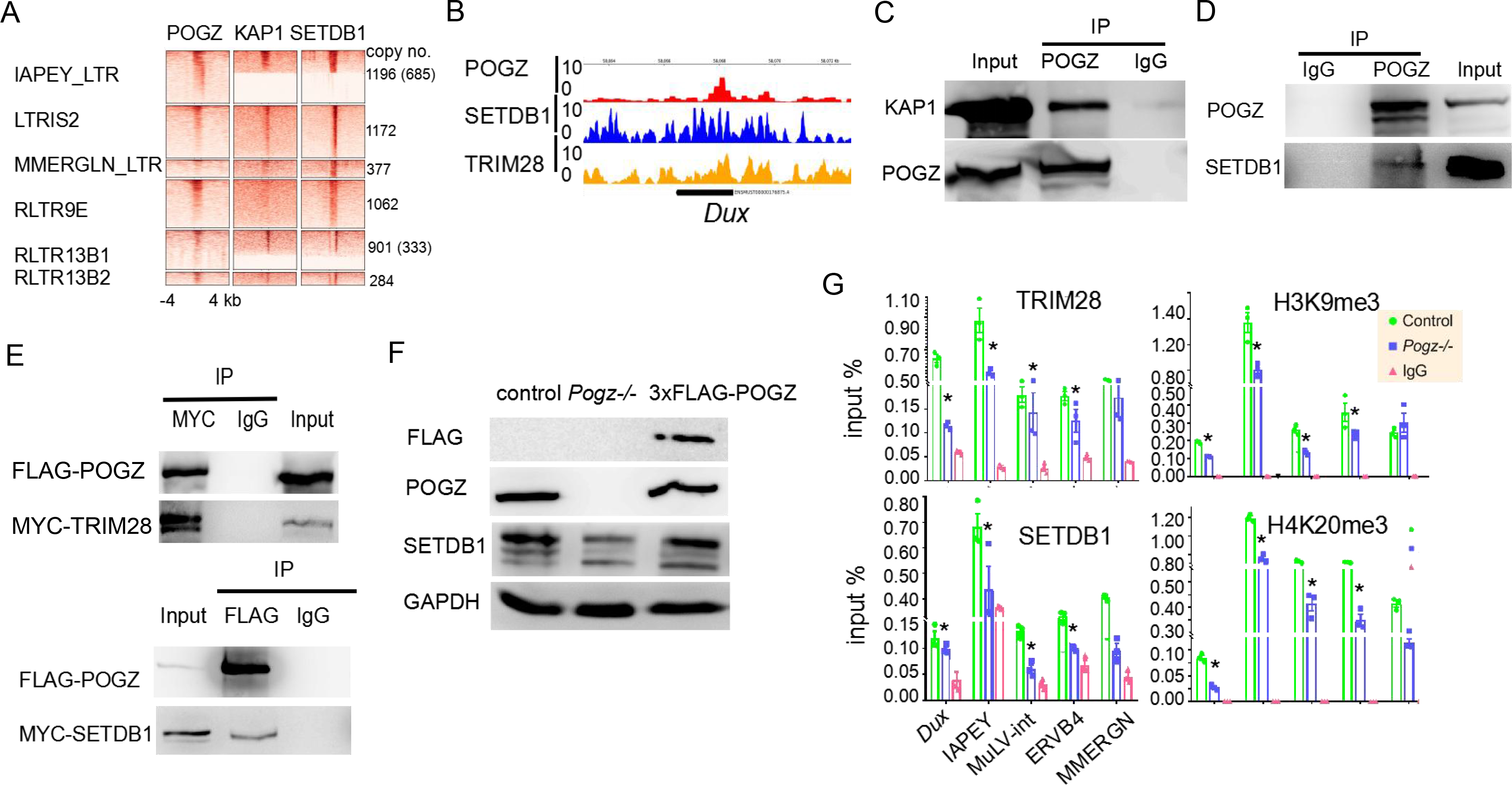
POGZ recruits TRIM28/SETDB1 to target ERVs. (A) Heat map of ChIP-seq data showing the enrichment of POGZ, SETDB1 and TRIM28 on the indicated ERVs in control and *Pogz−/−* ESCs. The ERV copy numbers were shown at right side. (B) Genomic view of ChIP-seq data for POGZ, SETDB1, TRIM28 on the *Dux* locus. (C) Western blot image showing that endogenous POGZ pulled down TRIM28/KAP1 in ESCs. (D) Western blot image showing that POGZ pulled down SETDB1 in ESCs. (E) Up: co-IP experiments showing Flag-POGZ interacts with MYC-TRIM28; Bottom: co-IP experiments showing Flag-POGZ associates with MYC-SETDB1. (F) Western blot image showing that SETDB1 levels were reduced in *Pogz−/−* ESCs, which can be rescued by reintroducing Flag-tagged POGZ. GAPDH was used as a loading control. (G) ChIP PCR results showing the enrichment of SETDB1, H3K9me3 H4K20me3, and TRIM28 on *Dux* and the indicated ERVs in control and *Pogz−/−* ESCs. WB and ChIP were repeated two times.

Next, we investigated whether POGZ interacts with TRIM28/SETDB1 by performing co-IP experiments. The results showed that endogenous POGZ can pull down both TRIM28 and SETDB1, albeit exhibiting more affinity for TRIM28 (Figure 6C-D). To further confirm this, we transfected MYC-tagged TRIM28 and FLAG-tagged POGZ into 293T cells, and performed the pull-down experiments. The results confirmed that MYC-tagged SETDB1 and FLAG-tagged POGZ can readily pull down each other (Figures 6E; Up). Similar results were observed for SETDB1 (Figures 6E; Bottom). Interestingly, we found that SETDB1 levels were slightly reduced in the absence of POGZ, which can be largely rescued by reintroducing the Flag-tagged POGZ (Figure 6F).

Next, we asked whether POGZ recruits the heterochromatin machinery containing TRIM28/SETDB1 to *Dux* and ERVs. To answer this question, we analyzed the chromatin status of POGZ-bound ERVs and the *Dux* gene. We found that there was a significant decrease of TRIM28, SETDB1, H3K9me3 and H4K20me3 levels on the selected ERVs and *Dux* in *Pogz−/−* ESCs (Figure 6G).

### Activation of POGZ-bound ERVs is associated with up-regulation of nearby genes

*POGZ* is one of the top risk genes that are frequently mutated in neurodevelopmental disorders (NDDs). It has been shown that de-repressed ERVs can act as alternative cis-elements (such as promoter or enhancers) to modulate the expression of neighboring genes (Karimi et al., 2011; Sundaram et al., 2017). It was therefore possible that deregulation of nearby genes due to aberrant expression of POGZ-bound ERVs may contribute to the disease pathology in *POGZ* patients.

To test this hypothesis in our ESC model, we first grouped genes based on their genomic distance from POGZ-bound ERVs, and analyzed the gene expression levels. We found that in the absence of POGZ, genes located closer to POGZ-bound ERVs such as RLTR9E were prone to be activated and exhibited significantly higher expression levels than genes far from the ERVs, analogous to that of MERVL (Figure 7A-B). Importantly, GO analysis of up-regulated DEGs that are located within 5-10 kb from the POGZ-bound ERVs revealed the enriched terms such as regulation of nervous system development and neuron differentiation, and regulation of synapse organization (Figure 7C), which suggested that POGZ depletion resulted in co-activation of RLTR9E and their nearby neural genes.

**Figure 7.**
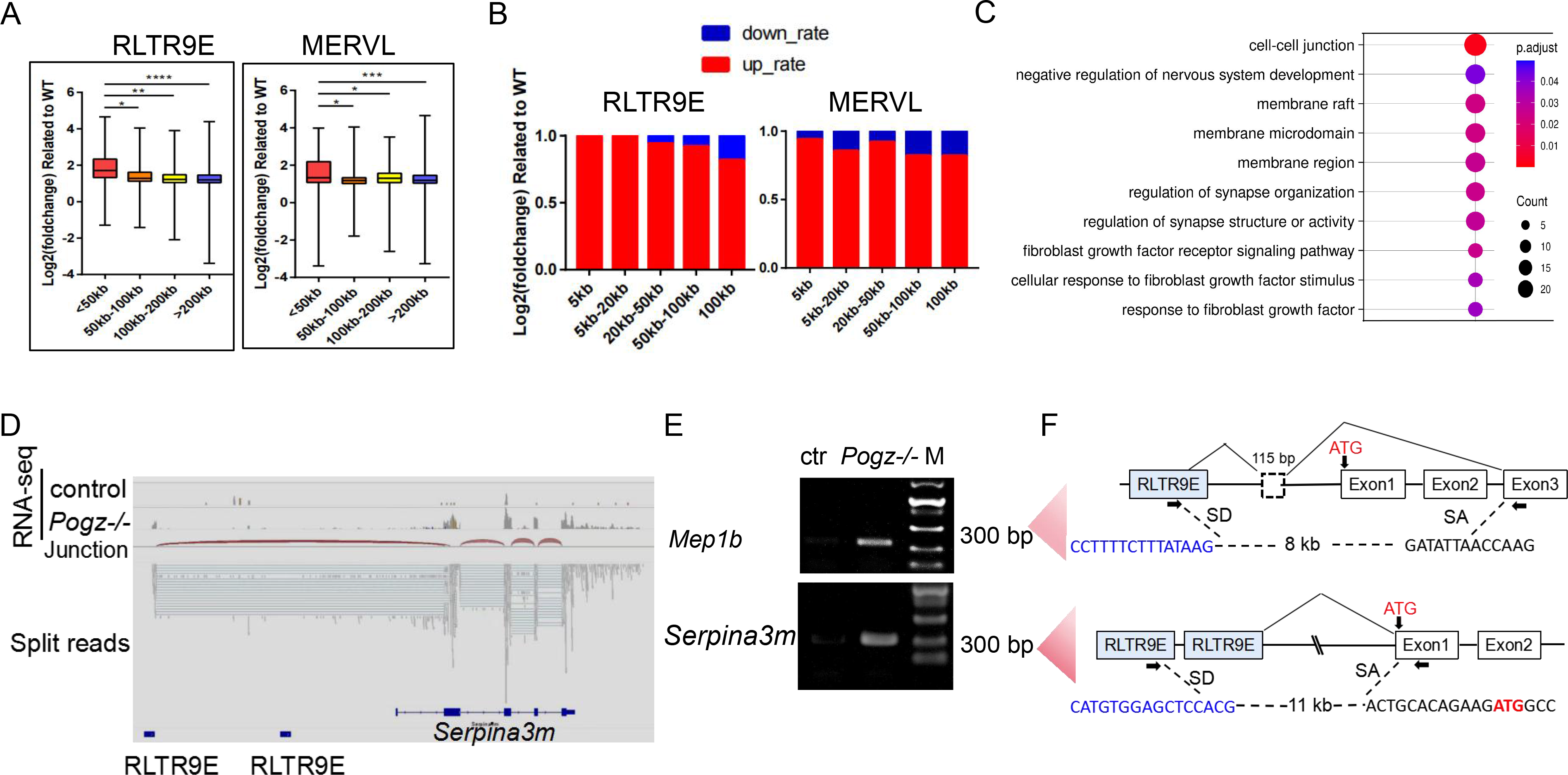
Increased expression of nearby genes is associated with derepression of POGZ-bound ERVs. (A) Graphs showing that in the absence of POGZ, genes located closer to POGZ-bound RLTR9E exhibited significantly higher expression levels than genes far from the ERV. MERVL was used as a positive control. (B) Proportion of up- and down-regulated genes in *Pogz−/−* ESCs is shown, according to their distance to the indicated REs. MERVL was used as a positive control. (C) GO analysis of up-regulated DEGs that are located within 5-10 kb from the POGZ-bound ERVs. (D) UCSC genome-browser screen shot of the 5’ end of the *Serpina3m* gene, showing RNA-seq tracks, alignment of the split paired-end RNA-seq reads in the locus, and RLTR9E upstream of the gene. (E) The presence of chimeric transcripts of the *Serpina3m* gene exclusively in *Pogz−/−* ESCs was validated by RT-PCR with primers (arrows in panel F) designed within the 50 bp regions to which the chimeric paired-end reads aligned. (F) Amplicons were cloned and sequenced, and the structure of the chimeric RNAs, the orientation and RLTR9E in which transcription initiates, and the annotated genic TSS and 50 exons (numbered) were shown for each gene. The sequence of the relevant novel splice donor (SD) and genic splice acceptor (SA) sites were also shown.

The *Serpina3m* gene was of particular interest to us for two reasons: 1) it was one of the most top genes that are activated by *Pogz* depletion (Supplementary Figure 1A-B), and 2) the *Serpina3* family genes have been shown to be significantly activated in mouse model of neural diseases or human disease tissues (Zhao et al., 2020; Kenigsbuch et al., 2022). We surveyed the paired-end RNA-seq reads for the presence of chimeric transcripts with one of the mate-pair reads aligning to RLTR9E and the other to annotated genic exons of *Serpina3m* (Figure 7D). To validate this, RT-PCR was performed using primers flanking RLTR9E and 5’ exons of *Serpina3m*. The results of gel electrophoresis showed that the PCR products were present in *Pogz−/−* ESCs but not in control ESCs (Figure 7E). The PCR products were then subjected to genotyping, which confirmed the formation of chimeric transcripts (Figure 7F; Supplementary Figure 7B-C). Similar results were observed for another gene *Mep1b* (Supplementary Figure 7A; Figure 7E-F). Taken together, we propose that aberrant expression of POGZ-bound ERVs may contribute to the disease pathology in *POGZ* patients by activating the expression of nearby neural genes.

## DISCUSSION

We have shown that POGZ is required for ESC cell survival and genome stability (Sun et al., 2022). However, the exact mechanisms remain unclear. Here, we showed that POGZ functions to maintain ESCs by directly targeting repetitive elements and *Dux*, and modulate H3K9me3/H4K20me3 marked heterochromatin by recruiting TRIM28/SETDB1 (Figure 8). The results were in line with previous reports that many of the same ERVs are de-repressed, and 2CLC state is induced in ESCs depletion of SETDB1 and TRIM28 (Rowe et al., 2010; Matsui et al, 2010; Wu et al., 2020). Also, our results showed that derepressed POGZ-bound ERVs can affect the expression of neighboring neural-related genes, which provides important insights into the disease pathology caused by POGZ dysfunction.

**Figure 8.**
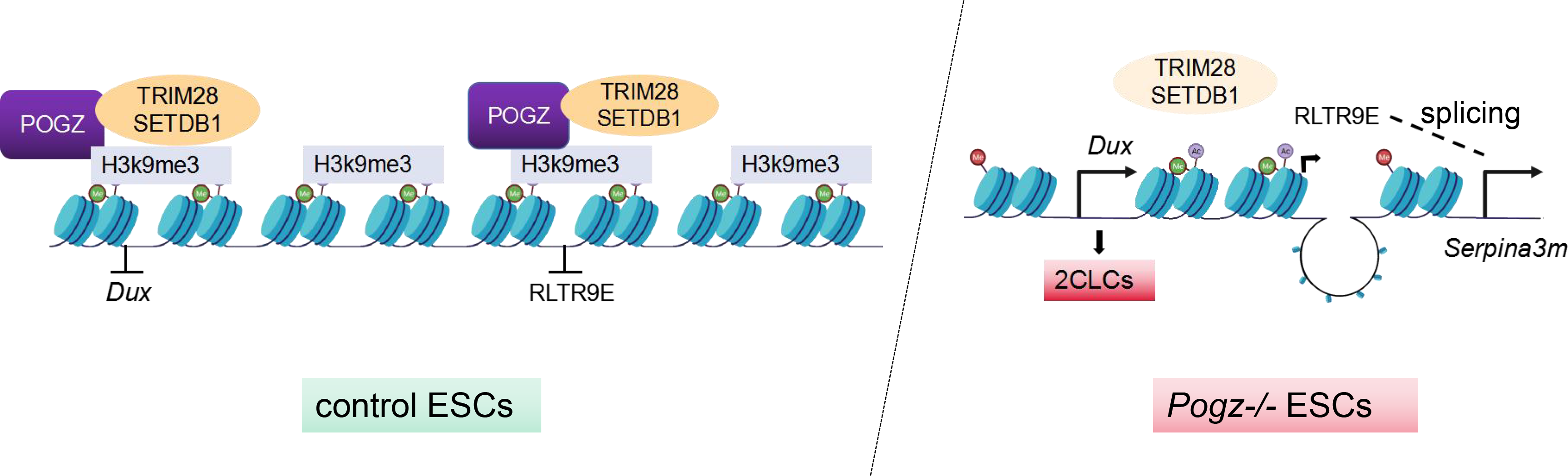
Cartoon illustrating the roles of POGZ in ESCs. Left: POGZ recruits TRIM28/SETDB1 to Dux and ERVs, thereby repressing their expression which is important for the maintenance of heterochromatin landscape and ESC identity. Right: in the absence of POGZ, H3K9me3 levels downstream of TRIM28/SETDB1 were reduced, which led to activation of *Dux* and POGZ-bound ERVs such as RLTR9E, resulting in transition to 2CLCs and genome instability. Chimeric transcripts formed by RLTR9E and its nearby genes such as *Serpina3m*, a gene up-regulated in neural disease condition.

*Dux* mRNAs are expressed at the early 2-cell stage in mice during the minor zygotic genome activation (ZGA) stage, whereas it activates downstream genes during major ZGA. Overexpression of Dux in mESCs results in changes in gene expression and endowed these cells with totipotency. Given its critical role in ZGA and ESCs to 2CLCs transition, the *Dux* gene must be tightly regulated. The upstream factors involved in transcriptional and epigenetic regulation of *Dux* have been identified, including developmental pluripotency associated 2/4 (DPPA2/4), Zscan4, LSD1, CAF1, and LIN28. In this work, we identified POGZ as a new regulator of the 2C-like state and ZGA gene transcription.

During early embryogenesis, different ERV sub-families are regulated in a highly stage-specific manner. In the field, an outstanding question is how different ERVs are targeted and repressed in physiological conditions. Although a few KRAB zinc-finger proteins such as ZFP809 and ZFP961 are identified for transcriptional silence of specific REs, other TFs remained to be determined (Imbeault et al., 2017; Tan et al., 2013; Wolf and Golf, 2009; Yang et al., 2022). In this work, we showed that POGZ binds to certain REs in a DNA sequence specific manner, analogous to KRAB zinc finger (KRAB-ZnF) proteins such as ZF809 and ZFP819. Previous work have shown that TRIM28, SETDB1, HP1, and the NuRD complex are members of a large repressive complex (Matsui et al., 2010; Ostapcuk et al., 2018). We propose that POGZ represses REs and *Dux* by recruiting a large repressive complex that includes TRIM28 and SETDB1.

Recent studies have suggested that REs might act as cis-elements capable of influencing expression of endogenous genes. Interestingly, the top POGZ-bound and regulated ERV families such as RLTR9E and RLTR13 were shown to be enriched for core pluripotency TF-binding sites (Sundaram et al., 2017). Furthermore, POGZ-bound TEs such as MuLV/RLTR4_Mm and RLTR6 can regulate host genes through modulating 3D genome structuring (Raviram et al., 2018; Wolf et al., 2015). These observations support our previous idea that POGZ may act as a ESC pluripotency-related factor as both a TF and a genome regulator (Sun et al., 2022).

The deregulation of REs is closely linked with human diseases, in particular neurodevelopmental disorders. For instance, ERV dys-regulation in the brain is thought to cause several neurodevelopmental and neurodegenerative disorders (NDDs) such as schizophrenia and amyotrophic lateral sclerosis (Douville et al. 2011), and LINE-1 activation drives ataxia (Takahashi et al., 2022). Of note, recent studies have emphasized the importance of differential splicing and isoform-level gene regulatory mechanisms in defining cell type and neural disease specificity (Gandal et al., 2018). *POGZ* is one of the top risk genes mutated in neurodevelopmental disorders (NDDs) such as ASD and ID. Our findings suggested that abnormal expression of POGZ-bound ERVs can contribute to the pathology of *POGZ*-related NDDs by affecting nearby neural genes. In the future, it will be interesting to further investigate the direct causal relationship in a mouse or primate model.

## Acknowledgments

This work was supported by the National Key Research and Development Program of China (2022YFA0806600), and National Natural Science Foundation of China (31671526) to YH Sun.

## Author contributions

XY Sun and LX Cheng performed the experiments; W Jiang, TZ Zhang and LX Cheng designed and performed bioinformatics analysis of the REs; YH Sun designed and provided the final support of the work, and wrote the paper. We thank Prof. He XM and Jiang H from Tongji Medical School of HUST for the help with the REs.

## Data availability

All RNA-seq, ATAC-seq, ChIP-seq and CUT&Tag data have been deposited in the public database at Beijing Genomic Institute (BGI) at https://bigd.big.ac.cn/, with the accession number of CRA003852. The *Pogz* mutant ESCs will be available to the community upon reasonable request.

## Ethics approval and consent to participate

This study was regulated and approved by the Institute of Hydrobiology, CAS, China.

## Consent for publication

Not applicable.

---

**Supplementary Figure 1.**
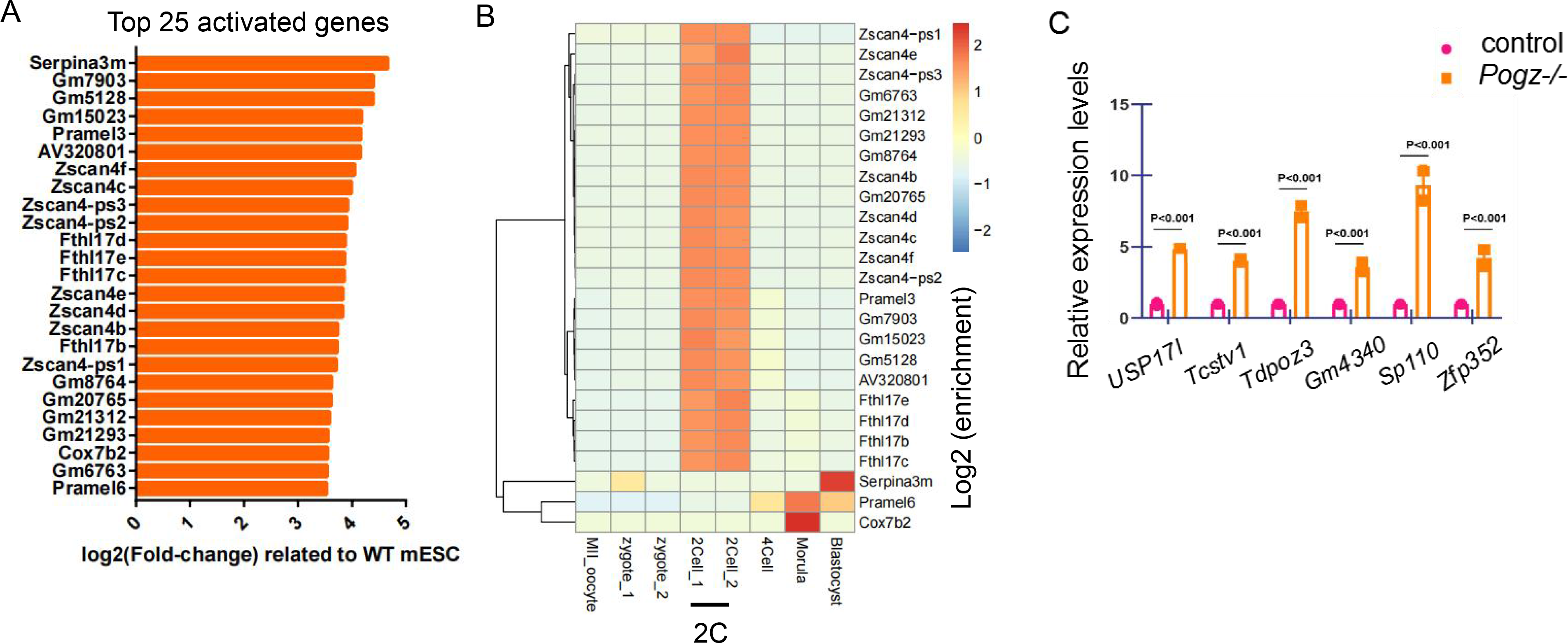
(A) Bar graph showing the top 25 activated genes in *Pogz−/−* ESCs. (B) Heat map showing that the top activated 25 genes were predominantly 2C-stage specific. (C) qRT-PCR analysis of selected 2C genes in control and *Pogz*−/− ESCs.

**Supplementary Figure 2.**
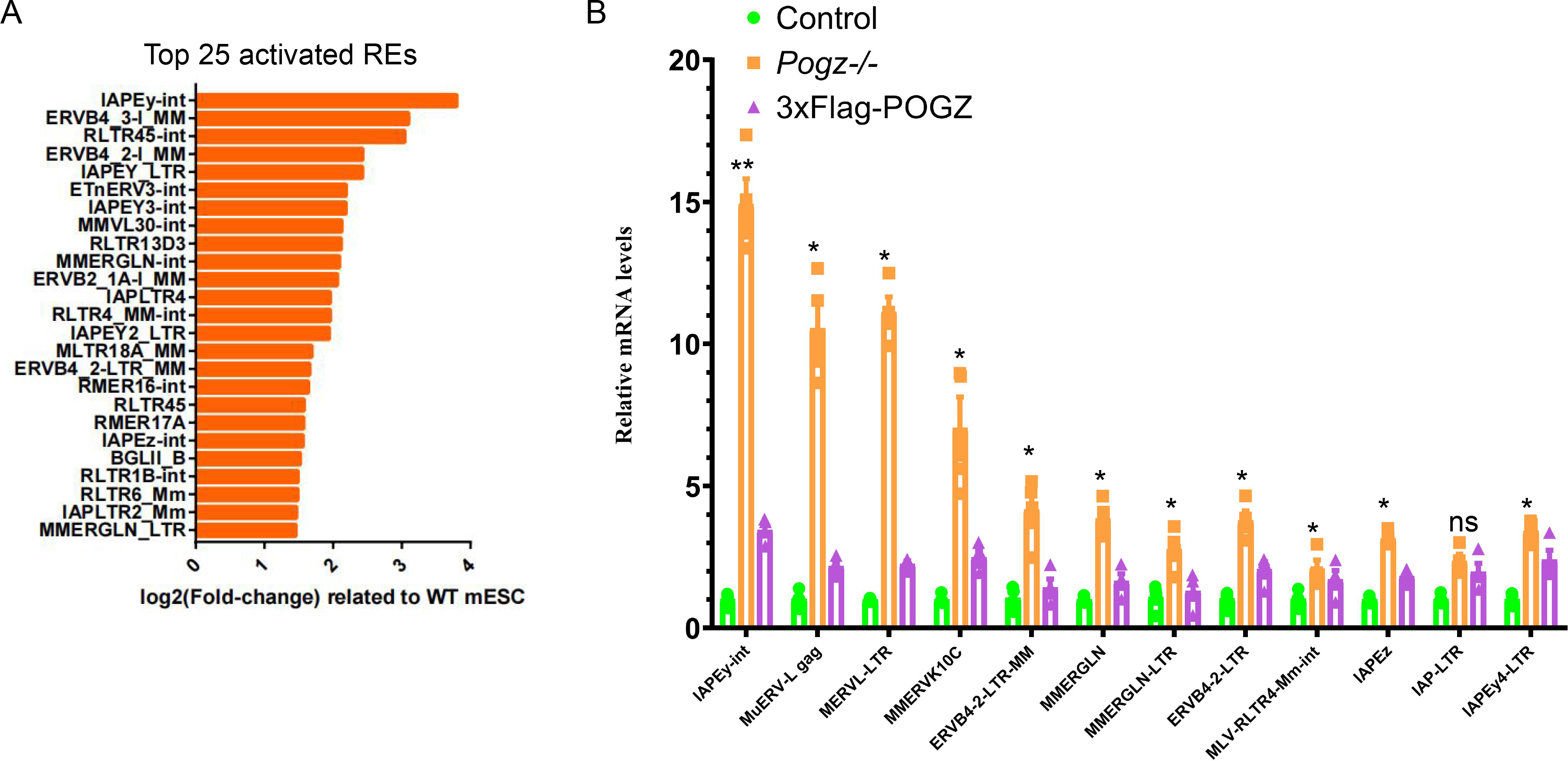
(A) Bar graph showing the top activated REs in the absence of POGZ. (B) The qRT-PCR analysis of selected ERVs in control, *Pogz*−/−, and Flag-POGZ restored *Pogz−/−* ESCs.

**Supplementary Figure 3.**
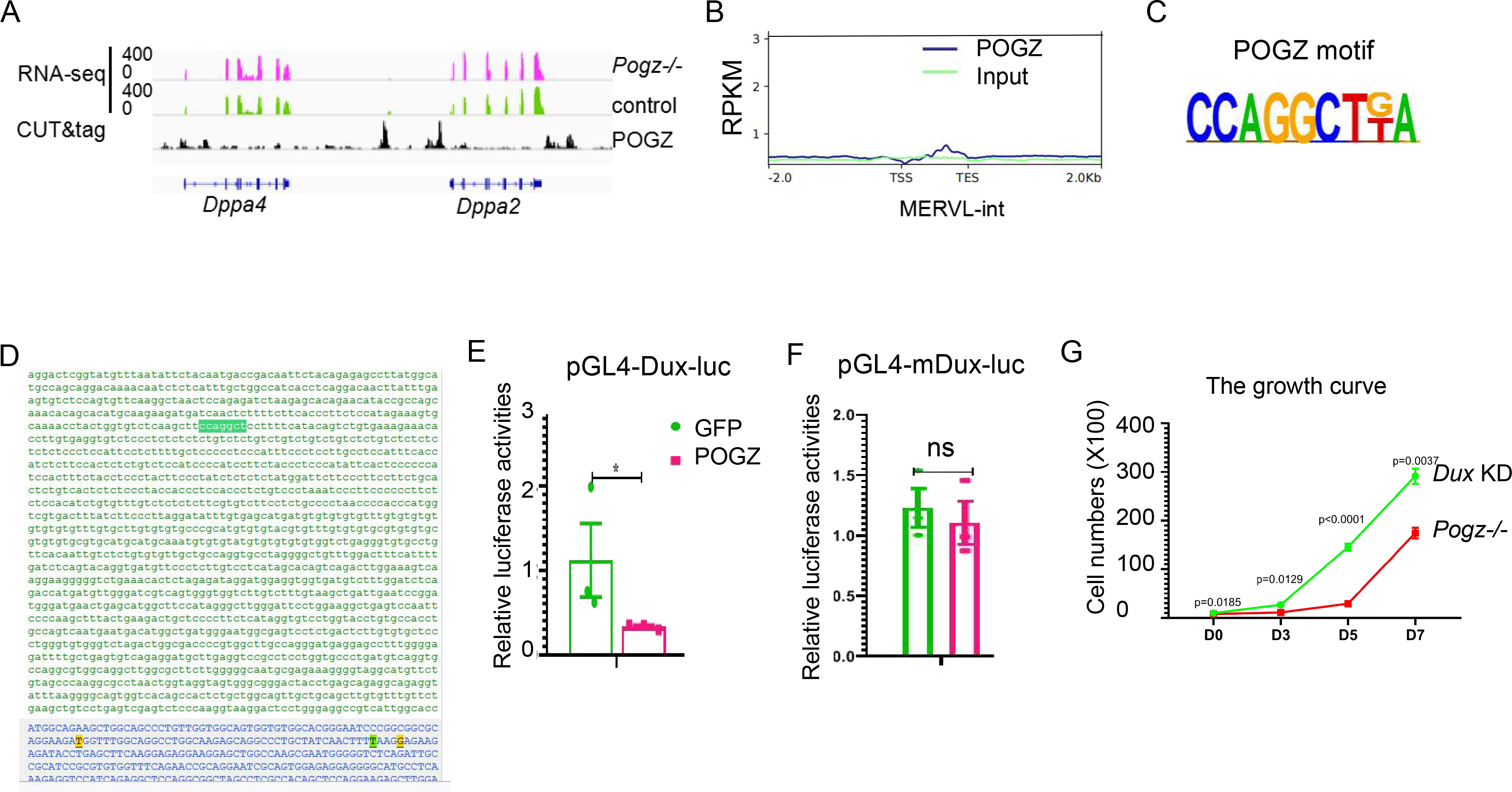
(A) Genomic views of RNA-seq and CUT&Tag data for the indicated genes. (B) Metaplot showing no POGZ enrichment on MERVL. (C) The mostly enriched POGZ binding motif based on CUT&Tag data. (D) DNA sequences of the *Dux* promoter, showing the position of the POGZ motif (highlighted). (E) Relative luciferase activities of pGL4-Dux-luc in the GFP and POGZ expressing 293T cells. (F) Relative luciferase activities of pGL4-mDux-luc in the GFP and POGZ expressing 293T cells. pGL4-mDux-luc is a mutated version of pGL4-Dux-luc, in which POGZ motif CCAGGCT was mutated to CCAaaCT. (G) *Dux* knockdown in *Pogz−/−* ESCs partially rescued the proliferation defects. The luciferase experiments were repeated two times.

**Supplementary Figure 4.**
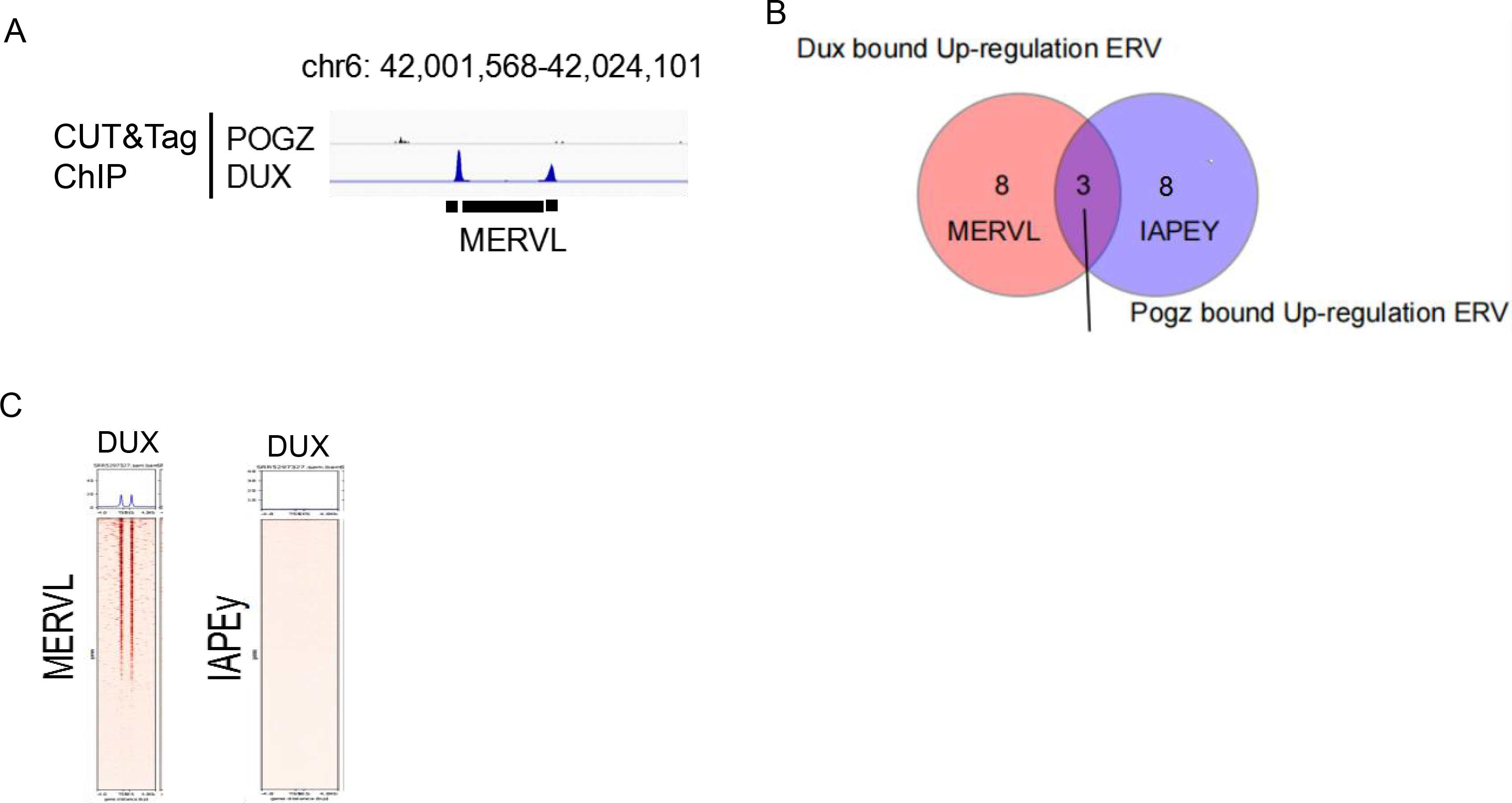
(A) Genomic view of MERVL locus on chromosome 6, showing that DUX but not POGZ was enriched. (B) Pie chart showing the overlap of DUX- and POGZ-bound up-regulated REs in its absence. (C) Heat map showing DUX occupancy on MERVL but not on POGZ-bound IAPEy, pointing that Dux does not directly regulate IAPEy.

**Supplementary Figure 5.**
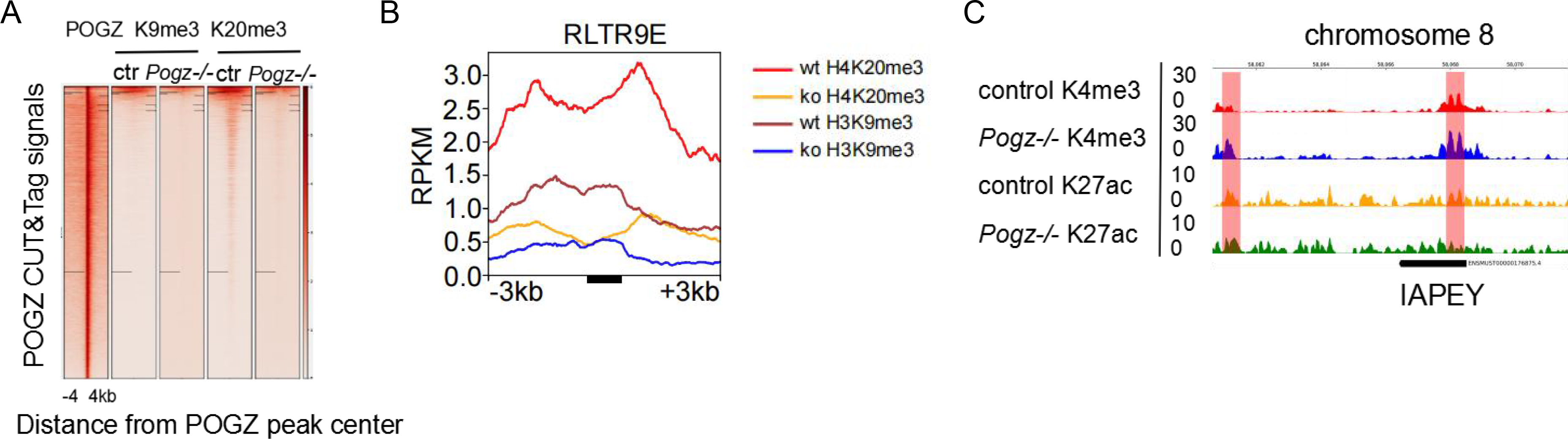
(A) Heat map showing the H3K9me3, and H4K20me3 ChIP-seq signals at POGZ binding sites in control and *Pogz−/−* ESCs. (B) Read count tag density pileups of the indicated ChIP profiles for POGZ-bound RLTR9E. RPKM: reads per kilobase of transcript, per million mapped reads. (C) Genomic view of ChIP-seq data for H3K4me3 and H3K27ac on POGZ-bound IAPEy in control and *Pogz−/−* ESCs.

**Supplementary Figure 6.**
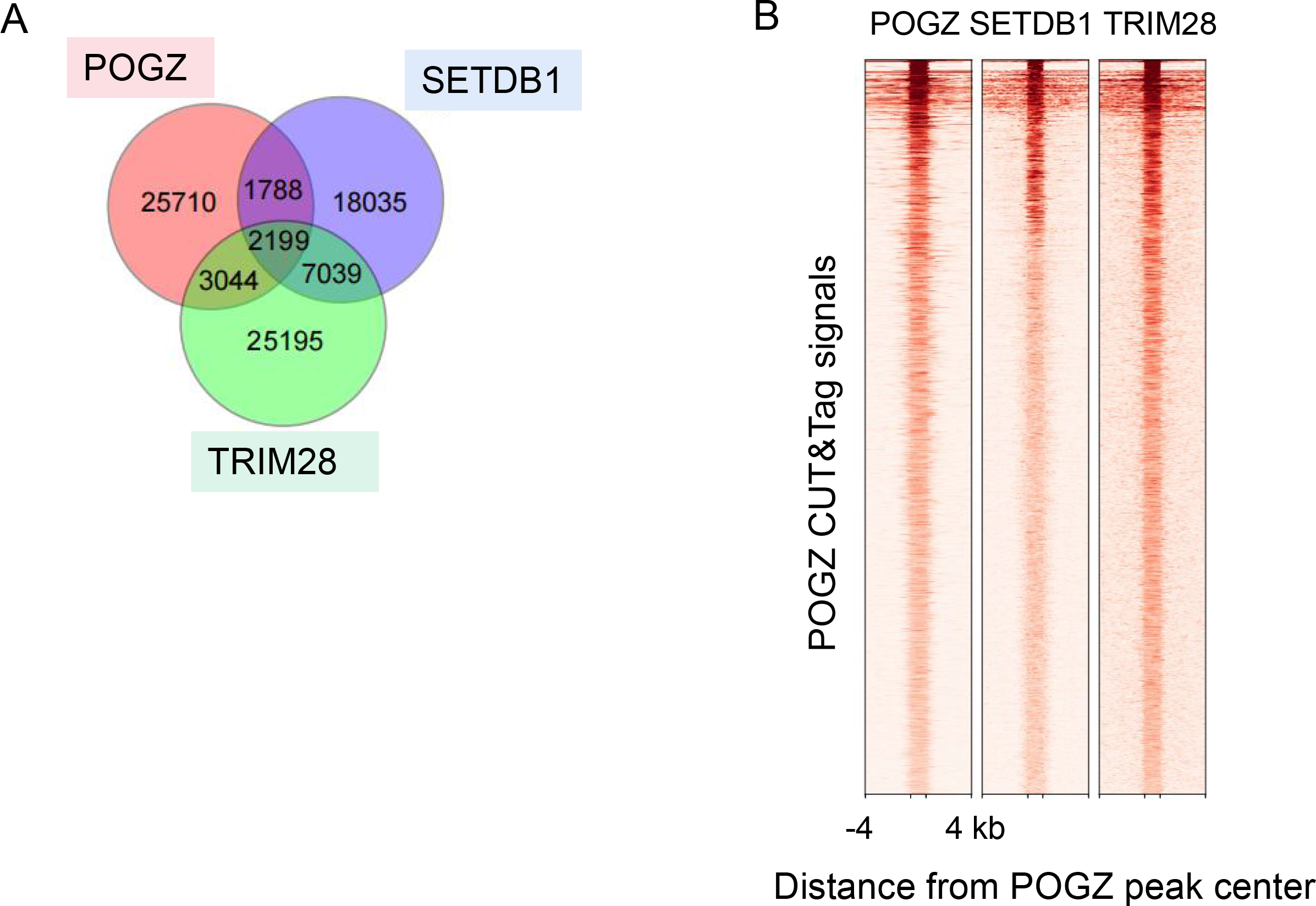
(A) Pie chart showing the co-localization of POGZ, SETDB1 and TRIM28 ChIP-seq peaks in ESCs. (B) Heat map showing the co-localization of POGZ, SETDB1 and TRIM28 genome-wide.

**Supplementary Figure 7.**
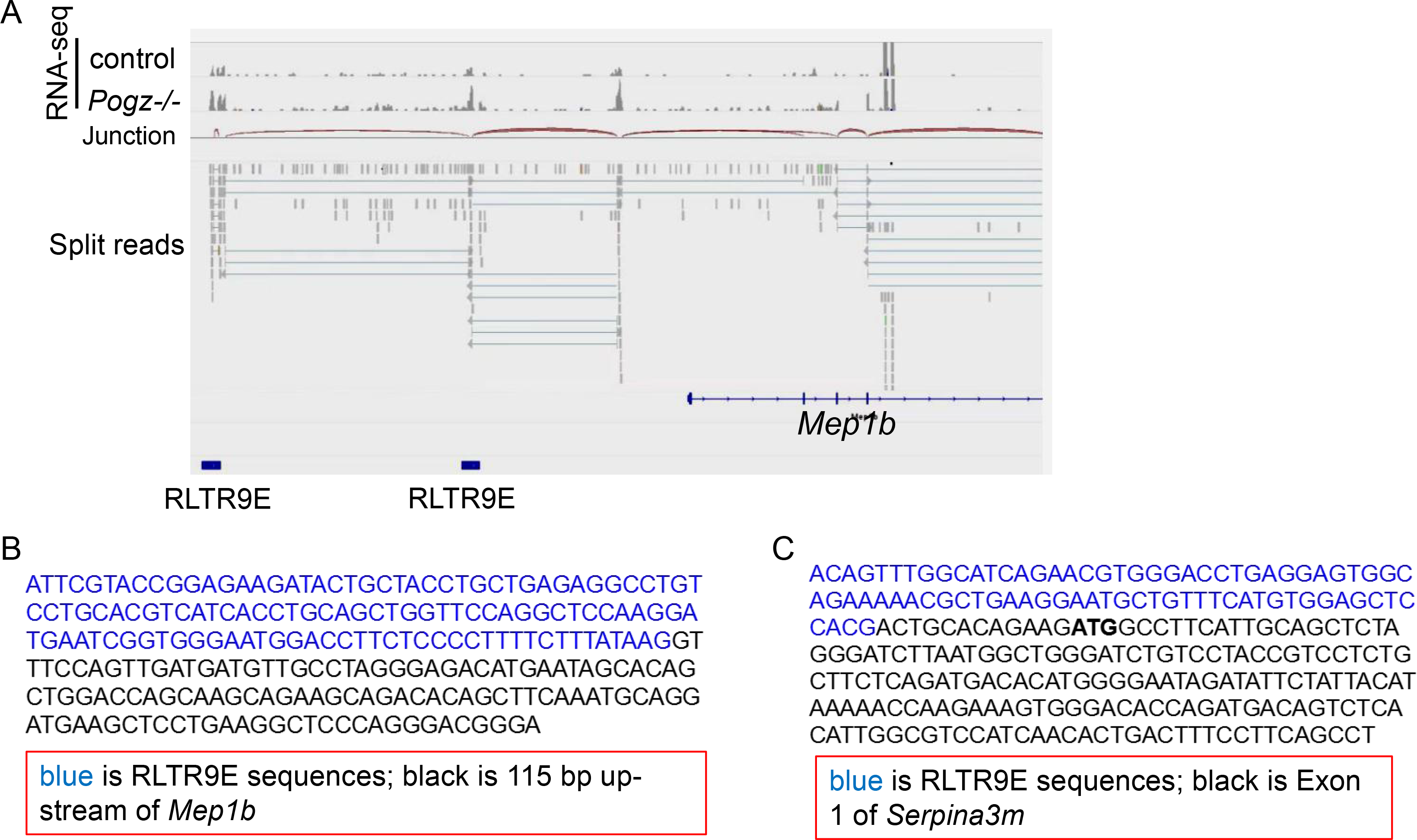
(A) UCSC genome-browser screen shot of the 5’ end of the *Mep1b* gene, showing RNA-seq tracks, alignment of the split paired-end RNA-seq reads in the locus, and RLTR9E upstream of the gene. (B) Genotyping results for PCR products of chimeric transcript (*Mep1b*) from Figure 7E-F. (C) Genotyping results for PCR products of chimeric transcript (*Serpina3m*) from Figure 7E-F.

## References

1. De Iaco, A., Planet, E., Coluccio, A., Verp, S., Duc, J., and Trono, D. (2017). DUX-family transcription factors regulate zygotic genome activation in placental mammals. Nat. Genet. 49, 941–945.

2. Douville R, Liu JK, Rothstein J and Nath A. (2011). Identification of Active Loci of a Human Endogenous Retrovirus in Neurons of Patients with Amyotrophic Lateral Sclerosis. ANN NEUROL, 69:141–151

3. Eckersley-Maslin, MA, Svensson V, Krueger C, Stubbs TM, Giehr P, Krueger F, Miragaia RJ, Kyriakopoulos C, Berrens RV, Milagre I, et al. (2016). MERVL/Zscan4 network activation results in transient genomewide DNA Demethylation of mESCs. Cell Rep. 17, 179–192.

4. Fukai R et al. (2015). A case of autism spectrum disorder arising from a de novo missense mutation in POGZ. J. Hum. Genet. 60, 277–279

5. Gandal et al. (2018). Transcriptome-wide isoform-level dysregulation in ASD, schizophrenia, and bipolar disorder. Science, 362, Issue 6420. DOI: 10.1126/science.aat8127

6. Gudmundsdottir, B. et al. (2018). POGZ is required for silencing mouse embryonic β-like hemoglobin and human fetal hemoglobin expression. Cell Rep 23, 3236–3248

7. Guo MY, Zhang YP, Zhou JF, Bi Y, Xu JQ, Xu C et al. (2019). Precise temporal regulation of Dux is important for embryo development. Cell Research 29: 956–959; https://doi.org/10.1038/s41422-019-0238-4

8. Hendrickson PG, Dora′ is JA, Grow EJ, Whiddon JL, Lim JW, Wike CL, Weaver BD, Pflueger C, Emery BR, Wilcox AL, et al. (2017). Conserved roles of mouse DUX and human DUX4 in activating cleavage-stage genes and MERVL/HERVL retrotransposons. Nat. Genet. 49, 925–934.

9. Hisada K, Sánchez C, Endo TA, Endoh M, Román-Trufero M, Sharif J, Koseki H, Vidal M. (2012). RYBP represses endogenous retroviruses and preimplantation- and germ line-specific genes in mouse embryonic stem cells. Mol Cell Biol 32: 1139–1149. doi:10.1128/MCB.06441-11

10. Hu ZH, Tan DE, Chia G, Tan HH, Leong HF, Chen BJ et al. (2020). Maternal factor NELFA drives a 2C-like state in mouse embryonic stem cells. Nat. Cell Biol., 22, 175–186

11. Iaco AD, Coudray A, Duc J, Trono D. (2019). DPPA2 and DPPA4 are necessary to establish a 2C-like state in mouse embryonic stem cells. EMBO Rep. 20, e47382

12. Ibaraki K et al. (2019). Expression analyses of POGZ, a responsible gene for neurodevelopmental disorders, during mouse brain development. Dev. Neurosci. 41, 139–148.

13. Ishiuchi T, Enriquez-Gasca R, Mizutani E, Boskovi A, Ziegler-Birling C, Rodriguez-Terrones D, Wakayama T, Vaquerizas JM, and Torres-Padilla ME. (2015). Early embryonic-like cells are induced by downregulating replication-dependent chromatin assembly. Nat. Struct. Mol. Biol. 22, 662–671.

14. Karimi M, Goyal P, Maksakova I, Bilenky M, Leung D, Tang JX, Shinkai Y, Mager D, Jones S, Hirst M, and Lorincz M. (2011). DNA Methylation and SETDB1/H3K9me3 Regulate Predominantly Distinct Sets of Genes, Retroelements, and Chimeric Transcripts in mESCs. Cell stem cell 8: 676–687.

15. Kaya-Okur et al., (2019). CUT&Tag for efficient epigenomic profiling of small samples and single cells. Nat. Commun. 10: 1930, https://doi.org/10.1038/s41467-019-09982-5

16. Kenigsbuch M, Bost P, Halevi S et al. (2022). A shared disease-associated oligodendrocyte signature among multiple CNS pathologies. Nat Neurosci 25, 876–886. https://doi.org/10.1038/s41593-022-01104-7

17. Li PS, Wang Li, Bennett B, Wang JJ, Li J, Qin YF, Takaku M, Wade PA, Wong JM and Hu G. (2017). Rif1 promotes a repressive chromatin state to safeguard against endogenous retrovirus activation. Nucleic Acids Research, 45, 12723–12738

18. Macfarlan et al. (2012). Embryonic stem cell potency fluctuates with endogenous retrovirus activity. Nature, 487: 57–64.

19. Macfarlan TS, Gifford WD, Agarwal S, Driscoll S, Lettieri K, Wang J, Andrews SE, Franco L, Rosenfeld MG, Ren B, and Pfaff SL. (2011). Endogenous retroviruses and neighboring genes are coordinately repressed by LSD1/KDM1A. Genes Dev. 25, 594–607.

20. Macfarlan TS, Gifford WD, Driscoll S, Lettieri K, Rowe HM, Bonanomi D, Firth A, Singer O, Trono D, and Pfaff SL. (2012). Embryonic stem cell potency fluctuates with endogenous retrovirus activity. Nature 487, 57–63.

21. Maksakova IA, Thompson PJ, Goyal P, Jones SJ, Singh PB, Karimi MM, Lorincz MC. (2013). Distinct roles of KAP1, HP1 and G9a/GLP in silencing of the two-cell-specific retrotransposon MERVL in mouse ES cells. Epigenetics Chromatin 6: 15. doi:10.1186/1756-8935-6-15

22. Markenscoff-Papadimitriou et al. (2021). Autism risk gene POGZ promotes chromatin accessibility and expression of clustered synaptic genes. Cell reports, 37: 110089

23. Matsumura et al. (2020). Pathogenic POGZ mutation causes impaired cortical development and reversible autism-like phenotypes. Nat. Commun. 11: 859. https://doi.org/10.1038/s41467-020-14697-z

24. Nozawa R et al. (2010). Human POGZ modulates dissociation of HP1alpha from mitotic chromosome arms through Aurora B activation. Nat. Cell Biol. 12, 719–727

25. Ostapcuk V. et al. (2018). Activity-dependent neuroprotective protein recruits HP1 and CHD4 to control lineage-specifying genes. Nature 557, 739–743

26. Payer L and Burns K. Transposable elements in human genetic disease. Nature Review Genetics. 20: 760 (2019)

27. Ran FA, Hsu PD, Wright J, Agarwala V, Scott DA, and Zhang F. (2013). Genome engineering using the CRISPR-Cas9 system. Nature Protocols 8, 2281–2308

28. Raviram et al. (2018). Analysis of 3D genomic interactions identifies candidate host genes that transposable elements potentially regulate. Genome Biology 19:216

29. Rowe HM et al. (2013). TRIM28 repression of retrotransposon-based enhancers is necessary to preserve transcriptional dynamics in embryonic stem cells. Genome Res. 23, 452–461

30. Rowe HM, et al. (2010). KAP1 controls endogenous retroviruses in embryonic stem cells. Nature 463, 237–240.

31. Ronan et al. (2013). From neural development to cognition: unexpected roles for chromatin. Nature Review Genetics, doi:10.1038/nrg3413

32. Stessman H. et al. (2016). Disruption of POGZ is associated with intellectual disability and autism spectrum disorders. Am. J. Hum. Genet. 98, 541–552

33. Stocking and Kozak. (2008). Murine endogenous retroviruses. Cell Mol Life Sci.; 65(21): 3383–3398

34. Suliman-Lavie R et al. (2020). *Pogz* deficiency leads to transcription dysregulation and impaired cerebellar activity underlying autism-like behavior in mice. NATURE COMMUNICATIONS 11: 5836 https://doi.org/10.1038/s41467-020-19577-0

35. Sun XY et al. (2020). ADNP promotes neural differentiation by modulating Wnt/β-catenin signaling. Nat. Commun. 11: 2984 https://doi.org/10.1038/s41467-020-16799-0

36. Sun XY, Cheng LX, Sun YH. (2022). Autism-associated protein POGZ controls ESCs and ESC neural induction by association with esBAF. Mol. Autism 13, 24

37. Sun Z et al. (2021). LIN28 coordinately promotes nucleolar/ribosomal functions and represses the 2C-like transcriptional program in pluripotent stem cells. Protein Cell.

38. Sundaram et al. (2020). Functional cis-regulatory modules encoded by mouse-specific endogenous retrovirus. NATURE COMMUNICATIONS | 8:14550 | DOI: 10.1038/ncomms14550

39. Tan B et al. (2016). A novel de novo POGZ mutation in a patient with intellectual disability. J. Hum. Genet. 61, 357–359

40. Takahashi et al. (2022). LINE-1 activation in the cerebellum drives ataxia. Neuron 110, 1–10

41. Vermeulen M. et al. (2010). Quantitative interaction proteomics and genome-wide profiling of epigenetic histone marks and their readers. Cell 142, 967–980.

42. Wang T et al. (2010). De novo genic mutations among a Chinese autism spectrum disorder cohort. NATURE COMMUNICATIONS 7:13316 DOI: 10.1038/ncomms13316

43. Whiddon JL, Langford AT, Wong CJ, Zhong JW, Tapscott SJ. (2017). Conservation and innovation in the DUX4-family gene network. Nat. Genet. 49, 935–940

44. White et al. (2016). POGZ truncating alleles cause syndromic intellectual disability. Genome Medicine 8: 3–11. DOI 10.1186/s13073-015-0253-0

45. Wu KX et al. (2020). SETDB1-mediated cell fate transition between 2C-like and pluripotent states. Cell Reports 30, 25–36

46. Yang B, Lu Fang, Gao QQ, Xu C, Xu JQ, Chen ZX, Wang YX, and Yang P. (2022). Species-specific KRAB-ZFPs function as repressors of retroviruses by targeting PBS regions. PNAS, 119 No. 11 e2119415119

47. Yang F, Huang X, Zang R, Chen J, Fidalgo M, Sanchez-Priego C, Yang J, Caichen A, Ma F, Macfarlan T et al. (2020). DUX-miR-344-ZMYM2-Mediated Activation of MERVL LTRs Induces a Totipotent 2C-like State. Cell Stem Cell, 26, 234–250.

48. Ye et al. (2015). De novo POGZ mutations are associated with neurodevelopmental disorders and microcephaly. Cold Spring Harb. Mol. Case Stud. doi:. 10.1101/mcs.a000455

49. Yu et al. (2021). rRNA biogenesis regulates mouse 2C-like state by 3D structure reorganization of peri-nucleolar heterochromatin. Nat. Commun. 12, 6365.

50. Yuan and Wessler. (2011). The catalytic domain of all eukaryotic cut-and-pastetransposase superfamilies. PNAS

51. Zalzman M et al. (2010). Zscan4 regulates telomere elongation and genomic stability in ES cells. Nature 464: 858–863. doi:10.1038/nature08882

52. Zhang et al. (2019). Zscan4c activates endogenous retrovirus MERVL and cleavage embryo genes. Nucleic Acids Res. 47, 8485–8501.

53. Zhao et al., (2019). Rare inherited missense variants of POGZ associate with autism risk and disrupt neuronal development. J. Genet. Genomics., 46: 247–257

54. Zhao et al. (2020). Alzheimer’s Risk Factors Age, APOE Genotype, and Sex Drive Distinct Molecular Pathways. Neuron 106, 727–742

